# C3-targeting siRNA Combined with Oseltamivir Alleviates Inflammation and Lung Injury in H1N1-Infected Mice

**DOI:** 10.64898/2026.11.16.766876

**Authors:** Gege Shi, Zixin Tu, Yu Li, Yawei Wang, Yu Ou, Shangqing Gao, Qi Ma, Xinyu Gao, Zimu Li, Yizheng Ao, Haoyuan Feng, Xiaofeng Han, Haitao Li, Minghui Yang, Xing-Jie Liang, Jinchao Zhang, Baoying Huang, Yuanyu Huang, Yuhua Weng

## Abstract

Influenza A H1N1 infection can trigger severe inflammatory responses that contribute to pulmonary tissue injury and disease progression. Aberrant complement activation is presumed to contribute to these pathological changes. Nevertheless, it remains unclear whether complement C3 serves as a shared upstream regulatory node linking inflammation and hemolytic injury. In this study, we established an H1N1-infected mouse model to dynamically monitor changes in inflammation, hemolysis and complement-related parameters throughout infection. We evaluated the C3-silencing efficacy of RG002, a liver-targeted siRNA against complement C3, as well as the therapeutic efficacy of RG002 combined with oseltamivir against H1N1 infection. Our results showed that H1N1 infection induced prominent inflammatory responses and alterations in peripheral blood cell parameters, accompanied by pulmonary histopathological injury in mice. Hepatic C3 expression and complement activity were simultaneously and markedly elevated. RG002 suppressed C3 expression in a dose-and time-dependent manner and reduced complement-mediated erythrocyte lysis in serum. In vivo therapeutic experiments demonstrated that on the basis of viral replication suppression achieved by oseltamivir, combination treatment with RG002 inhibited excessive complement activation. This combined regimen reduced pro-inflammatory cytokine levels, attenuated pulmonary histopathological injury, and improved selected peripheral blood cell parameters. Notably the low-dose RG002 combination group exhibited superior performance across multiple evaluation indices, suggesting that the degree of C3 suppression is preferable to influence therapeutic effucaty. Collectively, our findings implicate complement C3 in the inflammatory pathology associated with H1N1 infection and demonstrate the therapeutic potential of C3-targeting siRNA as an adjunct to antiviral treatment. This study provides experimental evidence supporting the combination of antiviral therapy and controlled complement modulation as a potential host-directed therapeutic strategy for severe influenza.

**Graphical Abstract:** H1N1 influenza virus infection triggers pulmonary inflammation and excessive complement activation, exacerbating lung injury. Here, we present a dual-targeted therapeutic strategy integrating antiviral treatment with host complement modulation. Oseltamivir suppresses viral replication, while RG002, a C3-targeting siRNA, silences hepatic C3 expression to attenuate complement overactivation. Their combined administration alleviates pulmonary inflammation and tissue injury, reduces proinflammatory cytokine levels, improves biochemical indicators, and decreases hepatic C3 expression and serum complement activity. This strategy offers a promising therapeutic approach for influenza-associated lung injury.

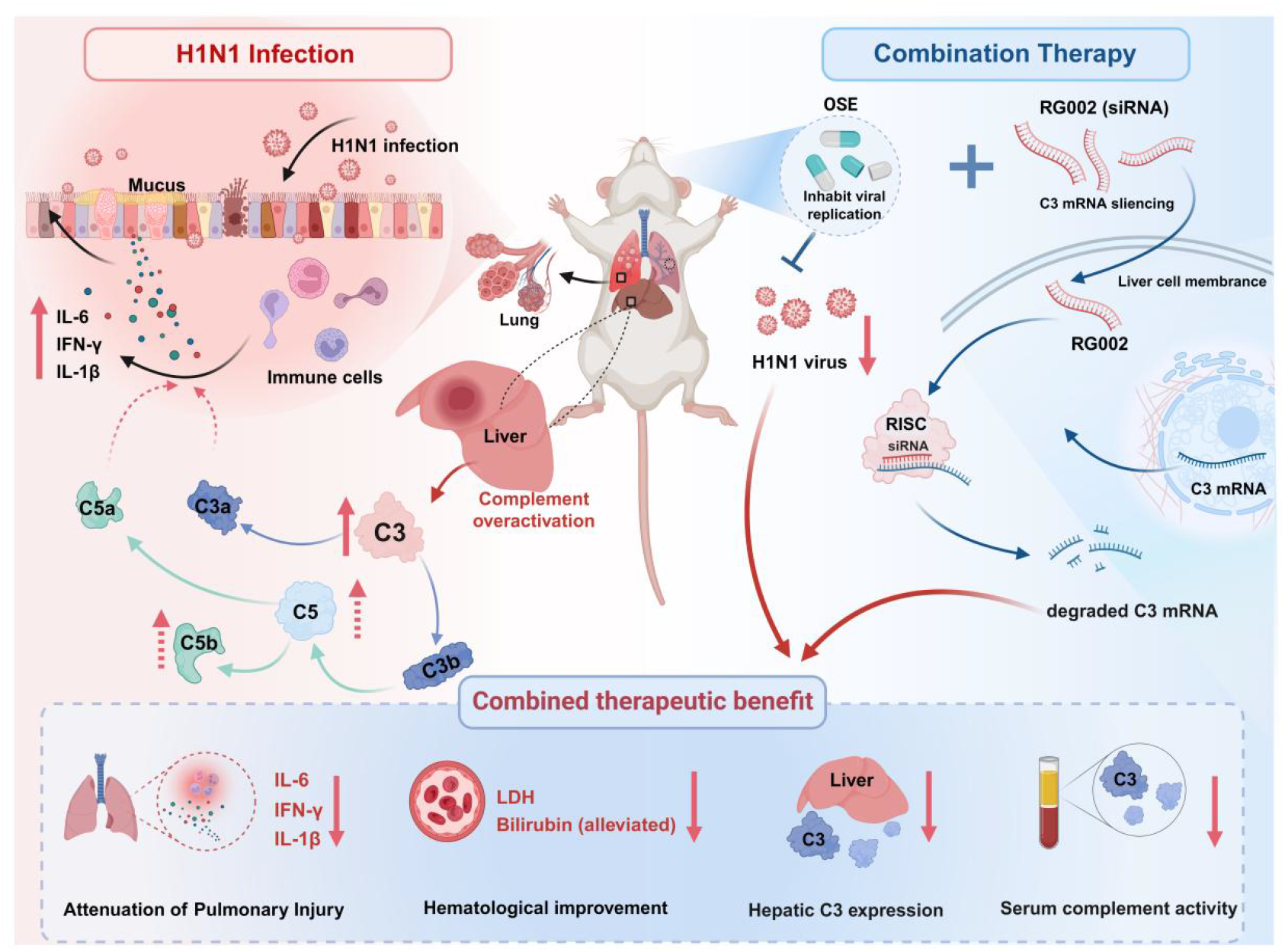

## Introduction

Influenza A virus (IAV) is a major cause of acute respiratory tract infections, with the H1N1 subtype continuing to pose a substantial threat to global public health(Cook et al., 2020; Taubenberger and Morens, 2020; Zakstelskaja et al., 1978). Current strategies for influenza prevention and treatment rely primarily on vaccination and antiviral agents, such as oseltamivir(Arevalo et al., 2022; Monto et al., 2025). However, the development and progression of severe influenza are driven not only by viral replication but also by dysregulated host immune responses(Nguyen et al., 2024). Excessive inflammation can lead to inflammatory cell infiltration, disruption of alveolar architecture, and pulmonary tissue injury, thereby exacerbating disease severity(Xiao et al., 2026). Therefore, developing adjunctive therapeutic strategies targeting excessive host inflammatory responses is of considerable importance for improving the treatment of severe influenza.

Following influenza virus infection, the host innate immune system recognizes viral components through pattern recognition receptors (PRRs)(Takeuchi and Akira, 2010), activating interferon signaling and downstream inflammatory pathways to restrict viral replication and spread. However, sustained or excessive immune activation can induce the release of large amounts of pro-inflammatory cytokines and chemokines, further aggravating pulmonary inflammation and tissue injury(Gu et al., 2021; Nguyen et al., 2024). In addition to local pulmonary inflammation, influenza virus infection may be accompanied by systemic pathological alterations, including changes in leukocyte subset distributions and erythrocyte-related parameters(Dengler et al., 2014). These hematological alterations may reflect complex systemic host responses to infection. Therefore, a comprehensive assessment of inflammatory markers, histopathological changes, and peripheral blood cell parameters may provide a more complete characterization of H1N1-associated pathological responses.

The complement system is an essential component of innate immunity and plays important roles in pathogen clearance, immune cell recruitment, and inflammatory regulation(Jayaraman et al., 2024; Mastellos et al., 2024). Appropriate complement activation contributes to antiviral defense through pathogen opsonization and the regulation of adaptive immune responses, whereas excessive or dysregulated activation may amplify inflammatory responses and contribute to host tissue injury(Xiao et al., 2024). Complement component C3 serves as the central convergence point of the classical, alternative, and lectin pathways and is a key mediator of complement amplification. Excessive C3 activation leads to the generation of the pro-inflammatory mediator C3a and promotes downstream C5a production, thereby contributing to inflammatory cell recruitment and pulmonary inflammation(Garcia et al., 2013; Song et al., 2018). Complement activation can also promote the formation of the terminal membrane attack complex (MAC), which may induce erythrocyte lysis under conditions of impaired complement regulation(Mastellos et al., 2024; Risitano et al., 2023). Notably, previous studies have shown that C3 deficiency may compromise host immune defense against influenza virus infection, suggesting that complete complement inhibition may not necessarily be beneficial for viral control(Kopf et al., 2002; O’Brien et al., 2011). Therefore, achieving a balance between preserving essential antiviral immune functions and limiting complement-mediated inflammatory injury represents an important consideration in the development of complement-targeted therapeutic strategies.

In recent years, complement-targeted interventions have emerged as promising approaches for modulating dysregulated immune responses(Mastellos et al., 2024). Compared with C5-targeting agents that act downstream in the complement cascade, targeting C3 offers the potential to regulate complement activity at a central convergence point, thereby limiting complement-associated inflammatory amplification(Ricklin, 2024). However, the protective role of C3 in antiviral immunity suggests that therapeutic outcomes may depend on the extent of complement inhibition. RNA interference (RNAi) technology provides a potential approach for targeted regulation of C3 expression(Jadhav et al., 2024; Tang and Khvorova, 2024). Small interfering RNA (siRNA) mediates sequence-specific degradation of target mRNA, thereby reducing the expression of the corresponding protein. As the liver is the primary source of circulating C3, liver-targeted siRNA-mediated C3 silencing represents a pharmacologically feasible strategy for modulating circulating complement levels and functional activity(Prado and Nagy, 2024; Zanchi et al., 2022). Furthermore, the magnitude and duration of siRNA-mediated C3 silencing can potentially be adjusted through dosing and treatment regimens, providing an opportunity to investigate the therapeutic effects of different degrees of complement suppression.

Antiviral therapy and host-directed immunomodulation target distinct aspects of influenza pathogenesis, providing a rationale for their combined application in severe influenza(Hui and Chan, 2025). Oseltamivir exerts its antiviral effects by inhibiting viral neuraminidase activity and limiting the release and spread of progeny virions(Pandey et al., 2023; Smith et al., 2011), whereas C3-targeting siRNA aims to modulate host complement activity and may alleviate inflammation associated with excessive complement activation(Zanchi et al., 2022). Combining these approaches may provide additional benefits by attenuating host inflammatory pathology while controlling viral replication(Pandey et al., 2023). However, whether siRNA-mediated C3 silencing can provide additional anti-inflammatory and tissue-protective effects when combined with oseltamivir in H1N1 infection remains to be determined.

In this study, we established an H1N1-infected mouse model to characterize infection-associated changes in inflammatory responses, complement activity, and peripheral blood cell profiles. We characterized the pharmacokinetic properties and C3-silencing efficacy, and evaluated its effects on serum complement-mediated hemolytic activity of RG002, a liver-targeted C3 siRNA. Furthermore, we investigated the therapeutic effects of two different doses of RG002 in combination with oseltamivir on inflammatory responses, pulmonary histopathological injury, and hematological alterations in H1N1-infected mice. Our study aimed to evaluate the therapeutic potential of C3-targeted siRNA as an adjunct to antiviral therapy and provide experimental evidence supporting host-directed therapeutic strategies for alleviating H1N1-associated inflammatory injury.

## Results

### H1N1 Infection Is Associated with Complement Activation, Pulmonary Inflammation, Airway Epithelial Injury, and Hematological Alterations

To investigate the effects of H1N1 infection on complement activation, pulmonary inflammation and injury, and hematological parameters, mice were infected with H1N1 virus at a dose of 3×LD₅₀. Lung tissues were collected on days 1, 3, 5 and 7 post-infection, and viral RNA levels were measured by RT-qPCR. Viral RNA was detected by day 1, reached high levels on days 3 and 5, and declined slightly on day 7 (Fig. 1A). No specific viral RNA signal was detected in uninfected controls. These findings indicate that viral RNA persisted in the lungs throughout the observation period.

**Figure 1.**
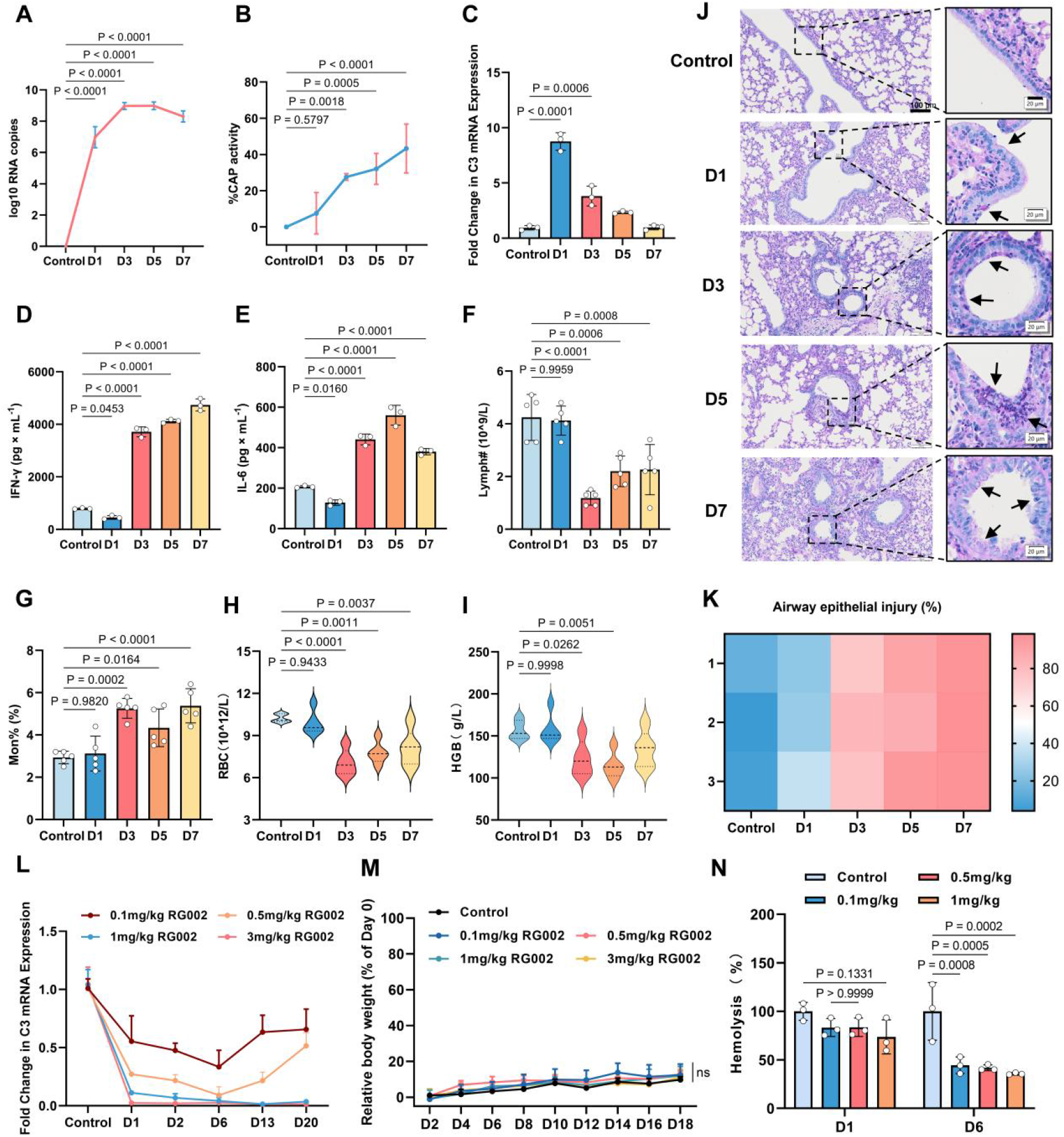
H1N1 infection-associated pathological alterations and pharmacodynamic charscterization of C3-targeting siRNA RG002 in mice. Mice were challenged with H1N1 influenza A virus at a dose of 3 LD₅₀, and samples were collected at 1, 3, 5 and 7 days post-infection to assess virological, immunological, hematological, and histopathological changes(A-K). Uninfected mice served as control. **(A)** Changes in viral RNA levels after H1N1 infection; **(B)** Serum alternation complement pathway activity. **(C)** Relative hepatic complement C3 mRNA expression; **(D, E)** Levels of pulmonary inflammatory cytokines IFN-γ and IL-6; **(F, G)** Changes in lymphocyte count (Lymph#) and monocyte percentage (Mon%); **(H, I)** Red blood cell count (RBC) and hemoglobin (HGB) levels **(J)** Representative periodic acid-Schiff (PAS)-stained lung sections showing airway epithelial injury and PAS-positive material accumulation following H1N1 infection. Black arrows indicate PAS-positive material and sloughed airway epithelial cells. Scale bars: 100 μm (main images) and 20 μm (magnified images); **(K)** Quantitative annalysis of the proportion of injured airway epithelium.

We next assessed whether H1N1 infection was associated with alterations in complement activity and C3 expression. Serum alternative complement pathway activity increased over the observation period, with a significant difference from uninfected controls observed from day 3 onward (Fig. 1B). We next examined hepatic C3 mRNA expression, as the liver is a major source of circulating complement proteins (Fig. 1C). C3 mRNA expression increased approximately 8.7-fold relative to the control level on day 1 and subsequently declined at later time points. Together, these findings show demonstrate distinct temporal patterns of hepatic C3 transcription and serum alternative complement pathway activity during H1N1 infection, characterized by an early increase in C3 mRNA expression and a progressive enhancement of complement activity.

To assess pulmonary inflammation following H1N1 infection, we measured interleukin-6 (IL-6) and interferon-γ (IFN-γ) levels in lung tissue homogenates using ELISA. Compared with uninfected controls, pulmonary IFN-γ levels in infected mice increased persistently starting on day 3 post-infection, and IFN-γ secretion surged to 4378 pg/mL on day 7 (Fig.1D). IL-6 concentrations were also markedly elevated following H1N1 infection, peaking at approximately 560.7 pg/mL on day 5. The temporal profile of IL-6 elevation largely paralleled changes in lung viral load, indicating prominent activation of antiviral immune responses during infection (Fig. 1E). These findings indicate that H1N1 infection induced pronounced pulmonary inflammatory responses during the middle and later stages of infection, with distinct temporal patterns of IFN-γand IL-6 production.

H1N1 infection also induced pronounced dynamic alterations in peripheral blood cell profiles. Lymphocyte counts were comparable to controls on day 1 but were significantly lower from days 3 through 7. In contrast, The proportion of monocytes increased markedly from day 3 and remained high in the late phase of infection. Red blood cell (RBC) counts dropped significantly on day 3 and showed a gradual recovery on days 5 and 7, although they remained below control levels. Hemoglobin (HGB) levels displayed a similar declining trend (Fig.1F-I). In summary, these findings demonstrate that H1N1 infection was accompanied by lymphopenia, an increased monocyte percentage, and reductions in erythrocyte-related parameters, indicating substantial hematological alterations during infection.

PAS staining revealed progressive histopathological alterations in the airway epithelium following H1N1 infection (Fig. 1J). PAS-positive material and early epithelial abnormalities were observed on day 1. More apparent epithelial disruption and focal detachment were observed from day 3 onward, accompanied by PAS-positive material and sloughed airway epithelial cells at later time points. Quantitative analysis revealed an increasing proportion of injured airway epithelium over the observation period, with the highest measured values observed on day 7 (Fig. 1K). Together, these results indicate that H1N1 infection was associated with increased complement activity, pronounced pulmonary inflammation, progressive airway epithelial injury, and hematological alterations.

### RG002 Induces Dose-and Time-Dependent Suppression of Hepatic C3 Expression

To characterize the magnitude and duration of RG002-mediated C3 silencing, uninfected mice received a single dose of RG002 at 0.1, 0.5, 1.0, or 3.0 mg/kg. Liver tissues were collected on days 1, 2, 6, 13, and 20 post-administration to determine hepatic C3 mRNA expression. RG002 induced a dose- and time-dependent reduction in hepatic C3 mRNA levels, with significant effects of dose, time, and their interaction (all P < 0.0001, two-way ANOVA; Fig. 1L). On day 1 post-administration, hepatic C3 mRNA expression was reduced by 45.1%, 73.0%, 88.7%, and 97.5% in mice receiving 0.1, 0.5, 1.0, and 3.0 mg/kg RG002, respectively, compared with untreated controls.

The magnitude and persistence of C3 silencing varied considerably across dose groups. At 0.1 mg/kg, RG002 produced relatively moderate C3 suppression, with a maximum reduction of 66.6%, followed by a gradual recovery in C3 mRNA expression from day 13 onward. Administration of 0.5 mg/kg RG002 resulted in more pronounced suppression, with C3 mRNA levels declining further during the early observation period before partially recovering at later time points. In contrast, treatment with 1.0 or 3.0 mg/kg RG002 induced rapid and sustained suppression of hepatic C3 expression, with mRNA levels remaining markedly reduced throughout the 20-day observation period (Fig. 1L). These findings demonstrate that RG002 effectively suppresses hepatic C3 expression over a broad dose range, with higher doses producing more profound and prolonged gene silencing.

To assess the effects of RG002 administration on body weight, mice were monitored throughout the observation period. Body weight increased gradually in all treatment groups, with no statistically significant differences in weight trajectories compared with controls (Fig. 1M). These observations indicate that RG002 administration was not associated with detectable adverse effects on body weight gain under the tested conditions.

### RG002 Reduces Serum Complement-Mediated Hemolytic Activity Ex Vivo

To determine whether RG002-mediated C3 silencing was accompanied by functional modulation of the complement system, serum complement-mediated hemolytic activity was evaluated using antibody-sensitized sheep red blood cells (SRBCs). Uninfected mice received RG002 at doses of 0.1, 0.5, or 1.0 mg/kg, and serum samples were collected on days 1 and 6 post-administration. The ability of serum samples to induce erythrocyte lysis was assessed by measuring absorbance at 542 nm. Serum from untreated control mice exhibited robust complement-mediated hemolytic activity, whereas serum collected from RG002-treated mice showed reduced hemolytic activity to varying degrees (Fig. 1N). On day 1 post-administration, hemolytic activity tended to decrease with increasing RG002 doses. However, none of the treated groups differed significantly from the control group, despite the substantial reductions in hepatic C3 mRNA expression observed at this time point (Fig. 1L, N). In contrast, on day 6, serum hemolytic activity was significantly reduced in all RG002-treated groups compared with controls. The greatest reduction was observed in the 1.0 mg/kg group, in which relative hemolytic activity decreased to approximately 35.7% of the control level (Fig. 1N).

Given the protective role of complement in antiviral immunity, we subsequently selected the 0.1 and 0.5 mg/kg doses of RG002 for combination treatment with oseltamivir in H1N1-infected mice. These doses exhibited distinct magnitudes and durations of C3 suppression, allowing us to investigate the therapeutic effects of different levels of C3-targeted intervention without employing the more profound and sustained silencing observed at higher doses.

In a separate experiment, uninfected mice were administered RG002 to evaluate its C3-silencing efficacy, effects on body weight and complement-modulating activity (L-N). **(L)**Relative hepatic C3 mRNA expression at different time-points following RG002 administration at doses of 0.1, 0.5, 1, and 3 mg/kg. Two-way ANOVA revealed significant effects of dose, time, and their interaction (all P < 0.0001); **(M)** Changes in body weight following administration of different RG002 doses. Two-way repeated-measures ANOVA revealed no significant differences in body-weight trajectories among treatment groups; **(N)** Serum complement-mediated hemolytic activity measured using antibody-sensitized sheep erythrocytes on days 1 and 6 following RG002 administration at doses of 0.1, 0.5, and 1 mg/kg. Data are expressed as mean ± SD, n = 3 mice per group. Ordinary one-way ANOVA was used for comparisons among multiple groups. Exact P values are shown in the graphs.

### RG002 Combined with Oseltamivir Improves Disease Outcomes in H1N1 - Infected Mice

Following confirmation of hepatic C3 suppression in vivo and inhibition of complement-mediated hemolysis in vitro, we further evaluated its therapeutic potential as monotherapy and in combination with the antiviral drug oseltamivir phosphate (OSE) in the H1N1 infection mouse model (Fig.2A). Survival and body-weight curve data showed that mice in the PBS vehicle group and the 0.5 mg/kg RG002 monotherapy group suffered rapid post-infection body-weight loss and all succumbed by day 7 post-infection. The 0.1 mg/kg RG002 monotherapy group and the OSE group yielded survival rates of 33.3% and 50.0%, respectively. In contrast, both combination-therapy groups achieved 100% survival and partially alleviated body-weight loss throughout the observation period, demonstrating greater protective efficay than either monotherapy (Fig. 2B, C).

**Figure 2.**
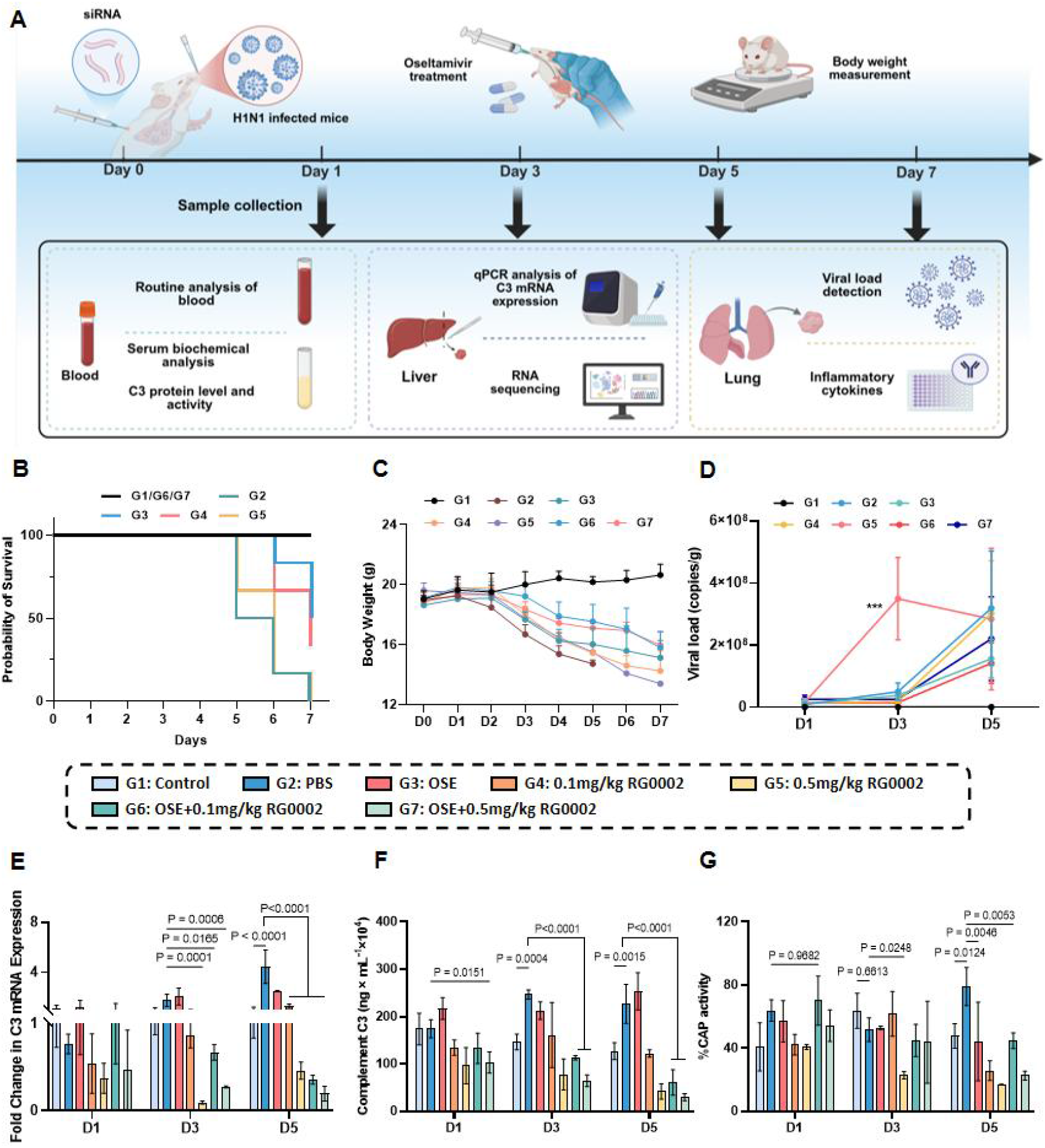
C3 RG002 combined with oseltamivir improves disease outcomes and effectively suppresses complement activation in H1N1-infected mice. **(A)** Schematic diagram of the therapeutic experimental design. Mice were challenged with H1N1 virus and administered RG002 on D0. oseltamivir treatment was initiated on the day of H1N1 challenge (D0) and administered daily thereafter. Blood, liver, and lung samples were collected at the indicated time points for subsequent analyses. Groups were defined as follows: G1, uninfected control; G2, H1N1-infected mice treated with PBS; G3, H1N1-infected mice treated with OSE; G4, H1N1-infected mice treated with 0.1 mg/kg RG002; G5, H1N1-infected mice treated with 0.5 mg/kg RG002; G6, H1N1-infected mice treated with OSE plus 0.1 mg/kg RG002; and G7, H1N1-infected mice treated with OSE plus 0.5 mg/kg RG002. **(B)** Survival curves of mice in each group after H1N1 infection. Mice reaching the predefined human endpoint of >25% loss of baseline body weight were recorded as non-survivors. **(C)** Body-weight dynamics of mice in each group during infection and treatment. **(D)** Measurement of H1N1 viral RNA loads in lung tissues. **(E)** Relative hepatic C3 mRNA measured by RT-qPCR at D1, D3, and D5. **(F)** Serum C3 concentrations measured by ELISA at D1, D3, and D5. **(G)** Serum complement alternative-pathway activity at D1, D3, and D5.. Data are presented as mean ± SD. Statistial analysis was performed using two-way ANOVA followed by multiple comparisons test compared with the PBS group. ***P < 0.001.

Pulmonary viral RNA levels were further quantified at days 1, 3, and 5 post-infection (Fig. 2D). No detectable viral RNA was observed in the uninfected control group, whereas PBS-treated mice showed a progressive increase in pulmonary viral burden over the courese of infection, reaching approximately 3 × 10^8^ copies/g on day 5.. Compared with the PBS vehicle group, the OSE group exhibited lower pulmonary viral burden, particularly at day 5, indicating that OSE effectively inhibits viral RNA levels and amplification in lung tissue. RG002 monotherapy differentially affected pulmonary viral burden depending on dose.. Viral loads in the 0.1 mg/kg RG002 group were largely comparable to those of the PBS vehicle group throughout the infection course, suggesting that low-dose C3 inhibition has limited influence on viral replication. However, the 0.5 mg/kg RG002 monotherapy group displayed elevated viral burden on days 3 and 5 post-infection, and showed significantly elevated pulmonary viral RNA levels on day 3 compared with the PBS group (P < 0.001).

Notably, in contrast to RG002 monotherapy, the OSE-containing combination groups showed lower pulmonary viral RNA levels. Between these groups, the OSE + 0.1 mg/kg RG002 combination group maintained the lowest viral burden on day 5 post-infection. These findings raise the possibility that combining antiviral treatment with C3 suppression may help control viral burden while limiting excessive complement activiaty.

To determine whether the treatment-associated differences in disease outcomes and pulmonary viral burden were accompanied by changs in the complement response, we next measured hepatic C3 mRNA expression-serum C3 concentrarion, and alternative-pathway activity (Fig. 2E-G).In the PBS-treated infected group, hepatic C3 mRNA expression increased across the sampled time points, with the highest observed level on day 5, the final sampling day (Fig. 2E). Meanwhile, serum C3 protein concentration was markedly elevated on day 5 (Fig. 2F), indicating that H1N1 infection promotes hepatic C3 transcription and increases circulating complement C3 levels. OSE treatment did not markedly alter the infection-associated changes in C3, as hepatic C3 mRNA expression and serum C3 concentrations showed overall profiles similar to those in PBS-treated mice. Monotherapy with 0.5 mg/kg RG002 significantly suppressed hepatic C3 mRNA up-regulation and the elevation of serum C3 protein, particularly at days 3 and 5, whereas the effects of 0.1 mg/kg RG002 were more modest.

Considering that H1N1 infection stimulates acute-phase complement synthesis in the liver, the C3-silencing efficiency of low-dose RG002 may be insufficient to counteract infection-driven C3 up-regulation, thereby maintaining relatively high C3 mRNA abundance. Notably, OSE + RG002 combination therapy further lowered hepatic C3 mRNA and serum C3 protein levels, with the OSE + 0.5 mg/kg RG002 group showing the most prominent inhibitory effect. Complement alternative-pathway (CAP) activity showed no significant difference between the treatment groups and the PBS group on day 1 post-infection. By day 3, CAP activity was significantly reduced in mice treated with 0.5 mg/kg RG002 compared with PBS-treated mice (P=0.0248), whereas no significant differences were observed for the other treatment groups. By day 5 post-infection, CAP activity was markedly increased in the PBS vehicle group, while all treatment groups exhibited significantly lower CAP activity relative to PBS. Among the treatment groups, 0.5 mg/kg RG002 monotherapy showed the lowest mean CAP activity (P<0.0001), and combination-therapy groups also kept activity at low levels (Fig. 2G). These results demonstrate that RG002 effectively supresses CAP activity during H1N1 infection, with a more pronounced effect at the higher dose.

Notably, OSE+RG002 combination regimens maintained lower hepatic C3 expression, circulating C3 levels, and CAP activity at later stages of infection, together with improved survival outcomes, indicating that the complement-modulatory activity of RG002 is preserved in the presence of OSE.

Previous results showed that combined treatment with RG002 and OSE improved survival outcomes in H1N1-infected mice while retaining modulatory effects on the complement pathway. To further determine whether these therapeutic effects were accompanied by attenuation of infection-associated inflammation, we subsequently assessed the dynamic changes in pulmonary pro-inflammatory cytokine levels and peripheral blood inflammatory cell populations.

ELISA analysis of lung homogenates showed that H1N1 infection induced a pronounced local inflammatory response. At day 1 post-infection, IL-6, IFN-γ, and interleukin-1β (IL-1β) levels were generally comparable among groups. By day 3, all three cytokines were markedly elevated in PBS-treated mice and remained high at day 5 (Fig. 3A-C). Compared with the PBS group, OSE treatment partially reduced inflammatory cytokine levels during the middle and late stages of infection. RG002/OSE combination therapy showed a more pronounced trend toward attenuation of the inflammatory response. In particular, the OSE + 0.1 mg/kg RG002 group maintained relatively low IL-6 and IFN-γ levels on both days 3 and 5, while the OSE + 0.5 mg/kg RG002 group also showed an inhibitory effect. IL-1β levels were markedly elevated in the PBS group during the middle and late stages of infection, whereas both combination regimens significantly reduced IL-1β levels, suggesting that complement modulation may be associated with attenuation of IL-1β-mediated inflammatory amplification. The inflammatory cytokine heatmap further illustrated these changes, with the low-dose combination group showing a more consistent low-cytokine profile (Fig. 3D). Overall, RG002 combined with OSE attenuated the accumulation of pro-inflammatory cytokines in the lungs of H1N1-infected mice, with the low-dose RG002/OSE combination exhibiting the most consistent control of the inflammatory response.

**Figure 3.**
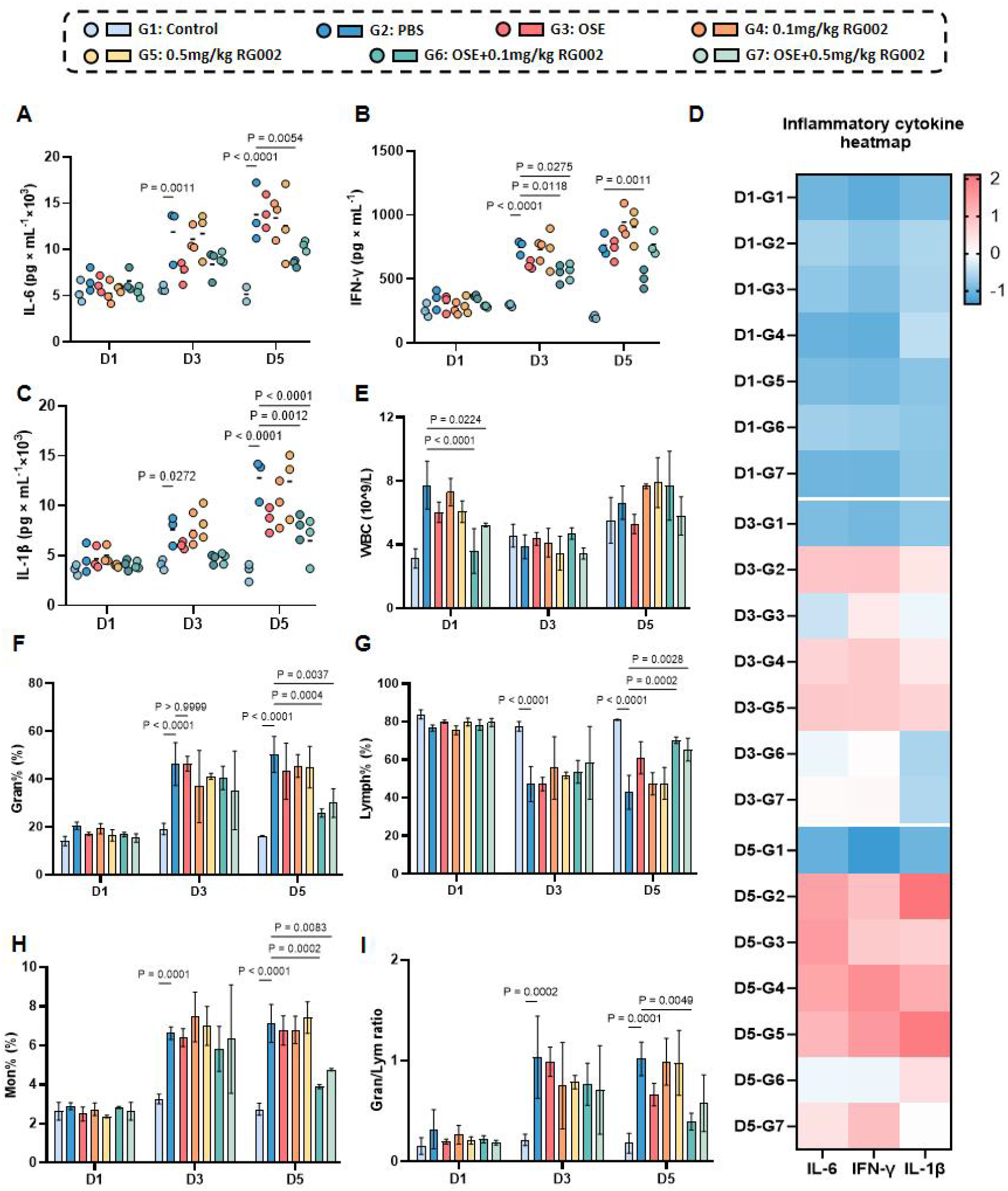
RG002 combined with oseltamivir alleviates pulmonary inflammation and improves peripheral leukocyte profiles in H1N1-infected mice. Experimental groups were defined as follows: G1, uninfected control; G2, H1N1-infected mice treated with PBS; G3, H1N1-infected mice treated with OSE; G4, H1N1-infected mice treated with 0.1 mg/kg RG002; G5, H1N1-infected mice treated with 0.5 mg/kg RG002; G6, H1N1-infected mice treated with OSE plus 0.1 mg/kg RG002; and G7, H1N1-infected mice treated with OSE plus 0.5 mg/kg RG002. **(A-C)** Pulmonary protein levels of IL-6, IFN-γ and IL-1β measured by ELISA. **(D)** Heatmap showing the relative levels of pulmonary inflammatory cytokines across treatment groups and time points. Date for each cytokine were individually Z-score normalized, with red indicating relatively higher levels and blue indicating relatively lower levels. **(E)** Peripheral-blood white blood cell count (WBC). **(F-H)** Percentages of granulocytes, lymphocytes, and monocytes in peripheral blood. **(I)** Granulocyte-to-lymphocyte ratio (Gran/Lymph ratio) in peripheral blood. Data are expressed as mean ± SD, n = 3 mice per group. Two-way ANOVA was applied for inter-group comparisons. P values for the indicated comparisons are shown in the graphs.

Peripheral-blood inflammatory-cell analysis revealed substantial alterations in leukocyte differential profiles upon H1N1 infection. Total white blood cell (WBC) counts increased sharply at early infection time points in the PBS vehicle group but total WBC levels subsequently fluctuated across groups. In comparison, changes in immune-cell subsets better reflect infection-elicited inflammatory conditions. Relative to the control group, the PBS vehicle group displayed elevated granulocyte percentage (Gran%), reduced lymphocyte percentage (Lymph%), and increased monocyte percentage (Mon%) in the mid-to-late infection phase, accompanied by a significantly higher Gran#/Lymph# ratio. Specifically on day 5, granulocyte counts were markedly elevated in the PBS vehicle group, whereas the OSE + 0.5 mg/kg RG002 combination group substantially lowered granulocyte counts, implying that combination therapy restrains excessive granulocyte-mediated inflammation at late infection stages. Furthermore, the OSE + 0.1 mg/kg RG002 combination group significantly increased lymphocyte counts and proportions and decreased the Gran#/Lymph# ratio as well as monocyte percentage on day 5, further confirming that RG002 combined with OSE rescues H1N1-induced peripheral immune-cell dysregulation (Fig.3E-I). Collectively, RG002 combined with OSE effectively alleviated infection-associated immune cell imbalance and dampens hyper-inflammation, consistent with its reduction of pro-inflammatory cytokine levels in lung tissue.

### RG002 Combined with Oseltamivir Modulates Serum LDH Activity and Bilirubin Profiles in H1N1-Infected Mice

Following the observation that the combination treatment attenuated pulmonary inflammatory responses and modulated peripheral leukocyte profiles, we further measured serum lactate dehydrogenase (LDH) activity and bilirubin concentrations to characterize infection- and treatment-associated biochemical alterations. Compared with the uninfected control group, serum LDH activity in the PBS-treated infected group showed no significant difference on day 1 but increased significantly on days 3 and 5 post-infection, consistent with increased cellular or tissue injury during infection. In contrast, the OSE group, 0.1 mg/kg RG002 monotherapy group and combination-therapy groups all reduced LDH levels to varying extents on days 3 and 5. The 0.1 mg/kg RG002 monotherapy group and OSE+0.5 mg/kg RG002 combination group achieved the significant improvement (Fig. 4A). Further bilirubin analysis revealed pronounced treatment-associated differences during the early stage of infection, without sustained elevations in bilirubin concentrations throughout the observation period. On day 1, total bilirubin (TBIL), direct bilirubin (DBIL) and indirect bilirubin (IBIL) levels were higher in PBS-treated infected mice than in uninfected controls (all P<0.0001). Notably, mice receiving the combination treatment maintained relatively lower levels of TBIL, DBIL, and IBIL on day 1 (Fig. 4B-D), suggesting that combination treatment was associated with favorable changes in selected serum biochemical parameters, particularly during the early stage of infection.

**Figure 4.**
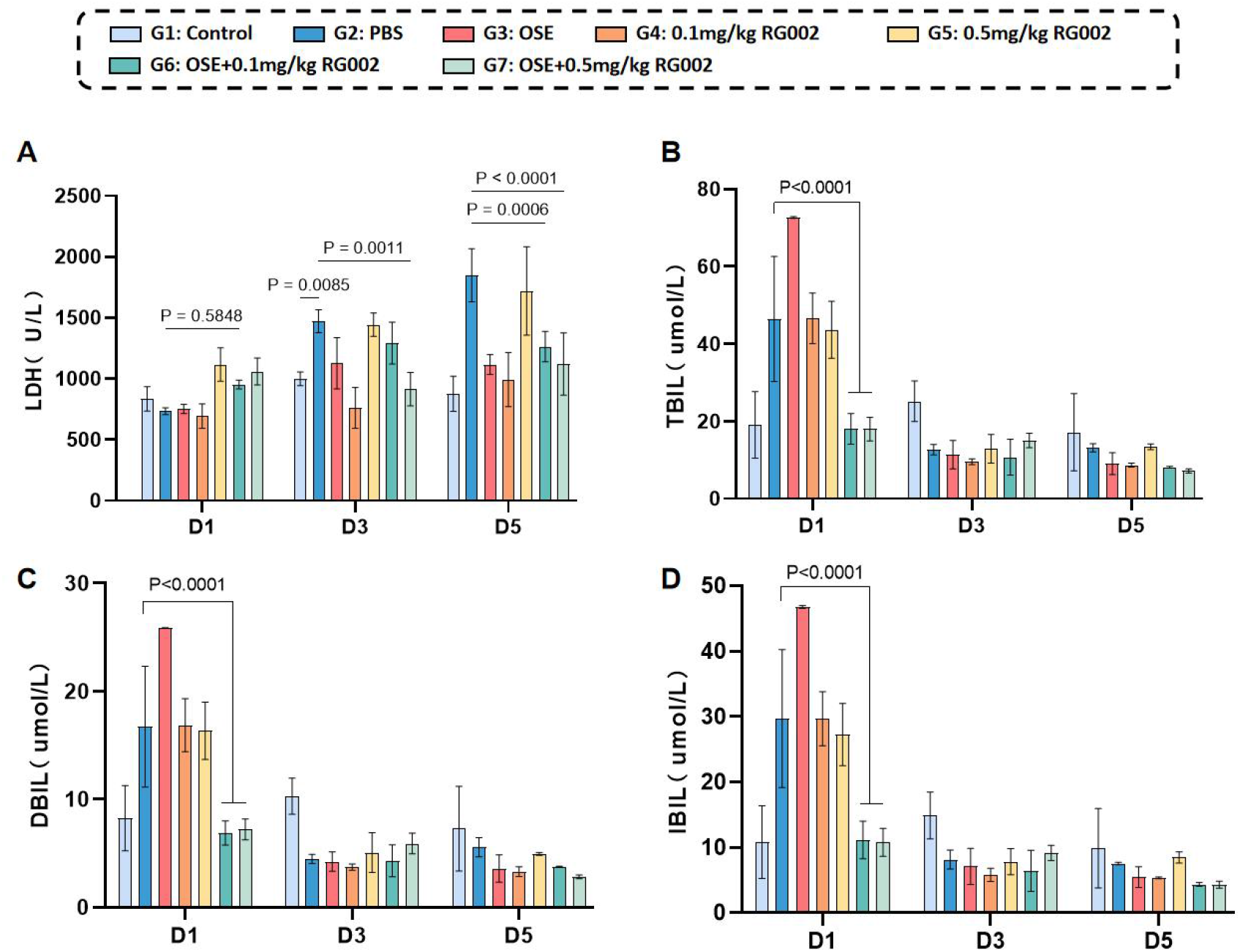
RG002 combined with OSE modulates serum biochemical parameters in H1N1-infected mice. Experimental groups were defined as follows:G1, uninfected control; G2, H1N1-infected mice treated with PBS; G3, H1N1-infected mice treated with OSE; G4, H1N1-infected mice treated with 0.1 mg/kg RG002; G5, H1N1-infected mice treated with 0.5 mg/kg RG002; G6, H1N1-infected mice treated with OSE plus 0.1 mg/kg RG002; and G7, H1N1-infected mice treated with OSE plus 0.5 mg/kg RG002. **(A)** Serum lactate dehydrogenase (LDH) levels; **(B)** Total bilirubin (TBIL) concentrations; **(C)** Direct bilirubin (DBIL) concentrations; **(D)** Indirect bilirubin (IBIL) concentrations. Data are expressed as mean ± SD, n = 3 mice per group. Statistical comparisons were performed using two-way ANOVA. P values for the indicated comparisons are shown in the graphs.

Overall, these findings indicate that H1N1 infection and subsequent treatments were accompanied by dynamic alterations in serum LDH activity and bilirubin profiles. Although RG002 combined with OSE was associated with improvements in selected biochemical parameters, these changes alone do not establish the occurrence or attenuation of hemolytic injury.

### RG002 Combined with OSE Alleviates H1N1-Induced Pulmonary Histopathological Injury

Hematoxylin and eosin (H&E) staining showed that H1N1 infection induced marked pulmonary pathological changes in the PBS group as early as Day 1, characterized by alveolar septal thickening, peribronchial inflammatory-cell infiltration, and focal disruption of alveolar architecture (Fig. S1A). These inflammatory and structural abnormalities were further aggravated by Day 5. In contrast, mice receiving combination therapy showed less inflammatory-cell infiltration and better-preserved pulmonary architecture (Fig. 5A). Quantitative analysis of inflammatory lesion areas in H&E-stained sections further showed that the PBS group had significantly larger inflammatory lesion areas than the Control group on both Day 1 and Day 5, whereas both low- and high-dose RG002 plus OSE combination treatments reduced the extent of inflammatory lesions (Fig. 5B; Fig. S1B), indicating that combined treatment with RG002 and OSE alleviates H1N1-induced pulmonary inflammatory injury.

**Figure 5.**
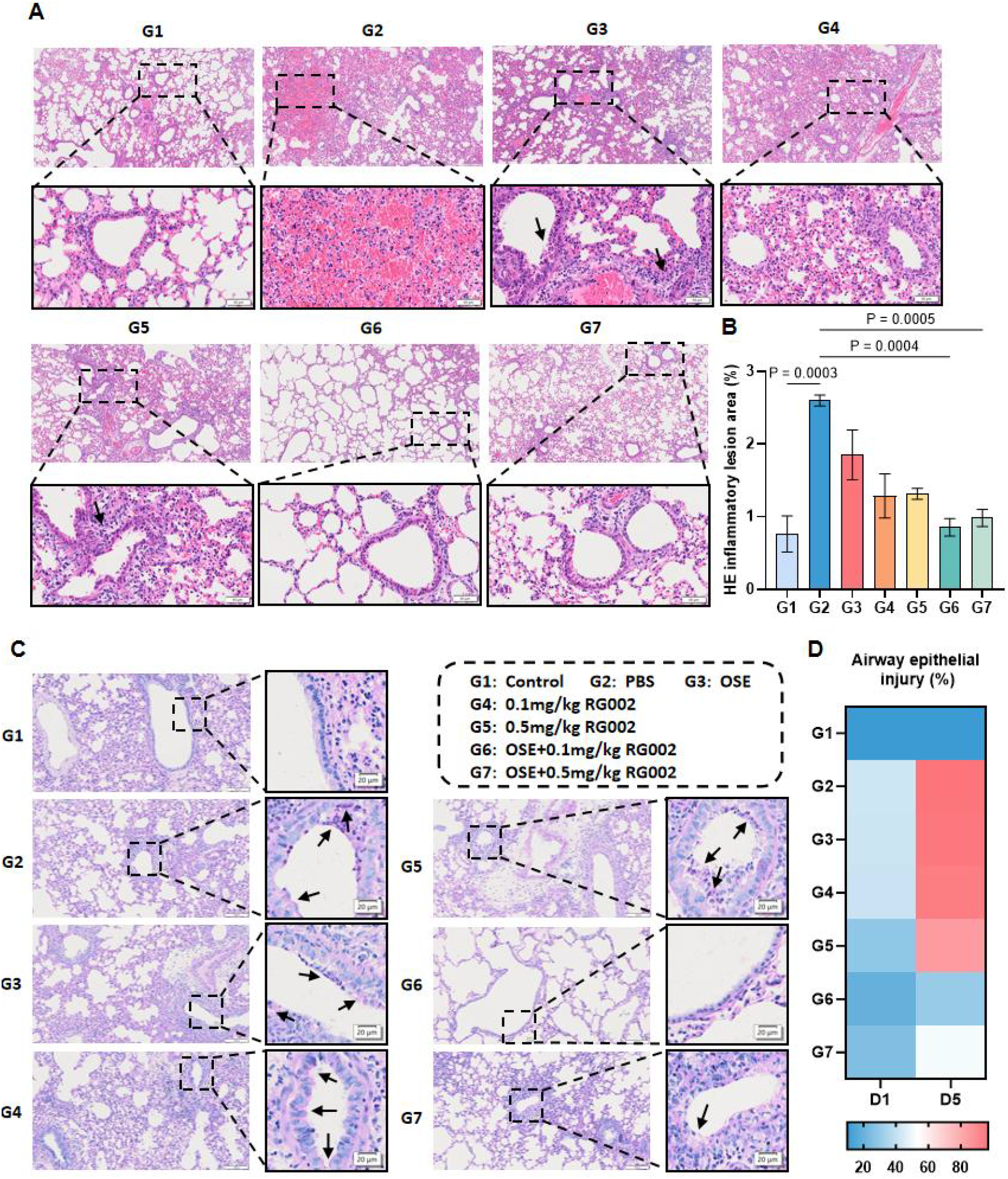
RG002 combined with OSE attenuates pulmonary inflammation and airway epithelial injury in H1N1-infection mice. Lung tissues collected from H1N1-infected mice under corresponding treatments were subjected to hematoxylin and eosin (H&E) and periodic acid-Schiff (PAS) staining. **(A)** Representative HE-stained images of lung tissues at day 5 post-infection, showing inflammatory cell infiltration and lung structural alterations upon infection and treatment. Black arrows denote areas of inflammatory cell aggregation or tissue damage. Scale bars: 200 μm (main panels), 50 μm (magnified insets). **(B)** Quantitative analysis of inflammatory lesion areas in HE-stained lung sections on day 5, used to evaluate the extent of pulmonary inflammatory injury. **(C)** Representative PAS-stained images of lung tissues at day 5, displaying airway epithelial structural integrity, epithelial cell shedding, and PAS-positive material deposition in the different treatment groups. Black arrows indicate PAS-positive material and shed airway epithelial cells. Scale bars: 100 μm (main panels), 20 μm (magnified insets). Data are expressed as mean ± SD, n = 3 mice per group. Ordinary one-way ANOVA was applied for inter-group comparisons. **(D)** Quantitative assessment of airway epithelial injury in PAS-stained lung sections on Days 1 and 5. Airway epithelial injury was defined as the percentage of the basement membrane length corresponding to injured airway epithelium relative to the total airway basement membrane circumference. The heatmap shows the percentage of injured airway epithelial circumference in each treatment group on Days 1 and 5, with colors ranging from blue to red indicating increasing severity of airway injury. Data are presented as mean ± SD, n = 3 mice per group. Statistical comparisons were performed using ordinary one-way ANOVA.

PAS staining further revealed marked histopathological abnormalities in the airway epithelium following H1N1 infection. On Day 1, prominent PAS-positive cells were observed in the airway epithelium of the PBS group, accompanied by apparent alterations in epithelial cell morphology. By Day 5, airway epithelial injury was further aggravated, as evidenced by extensive epithelial cell detachment, abnormal cellular morphology, and accumulation of PAS-positive material. Different degrees of airway epithelial injury remained detectable in the OSE and RG002 monotherapy groups. In contrast, the airway epithelial cells in the 0.1 mg/kg RG002 plus OSE combination group were relatively well organized and morphologically preserved, with no obvious epithelial cell detachment, suggesting a protective effect of low-dose combination treatment against infection-associated airway epithelial injury. The high-dose RG002 plus OSE combination group also showed partial preservation of airway epithelial architecture, although a certain degree of epithelial injury remained evident (Fig. 5C).We further quantified airway epithelial injury in PAS-stained sections. Using the airway basement membrane as a reference, the percentage of injured airway epithelial circumference was calculated as the length of the basement membrane corresponding to injured epithelium divided by the total airway basement membrane circumference. Quantitative analysis showed that airway epithelial injury increased over the course of H1N1 infection. In the PBS group, the percentage of airway epithelial injury was already elevated on Day 1 and increased further by Day 5. Mice receiving OSE or RG002 monotherapy still exhibited varying degrees of airway epithelial injury. In contrast, combination treatment with RG002 and OSE resulted in an overall reduction in airway epithelial injury on Day 5, with the low-dose combination group showing the most pronounced improvement, whereas the high-dose combination group also exhibited a partial reduction in epithelial injury (Fig. 5D).

### Hepatic transcriptomic analysis reveals attenuation of H1N1 infection-associated complement, coagulation, and inflammatory signaling pathways following combined RG002 and OSE treatment

To further r investigate the potential molecular mechanisms underlying the beneficial effects of combined RG002 and OSE treatment on H1N1 infection-associated inflammation and hematological abnormalities, hepatic transcriptomic analysis was performed on day 5 post-infection. Compared with the Control group, the PBS group exhibited marked alterations in gene expression profiles, with 676 upregulated and 567 downregulated genes identified (Fig. 6A). Among these, several genes associated with inflammatory signaling, complement regulation, and coagulation, including *Il1r1*, *Fga*, *Fgb*, *Fgg*, *C4b*, and *Cfh*, exhibited increased expression, suggesting that H1N1 infection induces transcriptional alterations in hepatic inflammatory responses and coagulation-related processes. To further evaluate the effects of combination therapy on hepatic gene expression, differential expression analyses were performed between the combination-treatment groups and the PBS group. A total of 338 upregulated and 399 downregulated genes were identified in the OSE plus 0.1 mg/kg RG002 group, whereas 134 upregulated and 122 downregulated genes were identified in the OSE plus 0.5 mg/kg RG002 group (Fig. 6B–C). Notably, both combination regimens were associated with marked downregulation of the target gene *C3* and the inflammation-related gene *Il1r1*. Several genes involved in coagulation and acute-phase responses, including *Fga*, *Fgb*, *Fgg*, *Thbd*, and *Serpine1*, also exhibited varying degrees of reduced expression. In addition, several inflammation-associated genes, including *Cxcl1*, *Saa1*, *Saa2*, and *Lcn2*, were downregulated in the lower-dose combination group (Fig. 6B–C). Collectively, both combination regimens were associated with alterations in hepatic gene expression profiles, with a greater number of differentially expressed genes observed in the lower-dose combination group. These transcriptional changes involved biological processes related to complement regulation, inflammatory responses, and coagulation. To systematically evaluate the biological pathways associated with H1N1 infection and therapeutic intervention, gene set enrichment analysis (GSEA) was performed. The results showed that H1N1 infection was associated with significant positive enrichment of multiple complement-, inflammation-, and coagulation-related pathways, including complement and coagulation cascades, TNF signaling, acute inflammatory response, IL-1 signaling, Toll-like receptor cascades, fibrin clot formation, and platelet activation-related pathways (Fig. 6D). Notably, the complement and coagulation cascades pathway was significantly positively enriched in the PBS group compared with the Control group (normalized enrichment score [NES] = 2.14, false discovery rate [FDR] = 5.03 × 10⁻⁶), suggesting enhanced hepatic transcriptional responses associated with complement and coagulation following H1N1 infection. Compared with the PBS group, the OSE plus 0.1 mg/kg RG002 group exhibited significant negative enrichment of all the selected pathways (FDR < 0.05). Similarly, most complement-, inflammation-, and coagulation-related pathways, including complement and coagulation cascades, IL-1-related signaling, Toll-like receptor cascades, and acute inflammatory response, were significantly negatively enriched in the OSE plus 0.5 mg/kg RG002 group. However, the NOD-like receptor signaling pathway remained positively enriched, whereas the negative enrichment of the TNF signaling pathway did not reach statistical significance (Fig. 6D). In particular, the complement and coagulation cascades pathway exhibited significant negative enrichment in both combination-treatment groups (Fig. 6E). These findings suggest that combined RG002 and OSE treatment is associated with an overall attenuation of H1N1 infection-induced hepatic transcriptional responses involving complement, inflammation, and coagulation.

**Figure 6.**
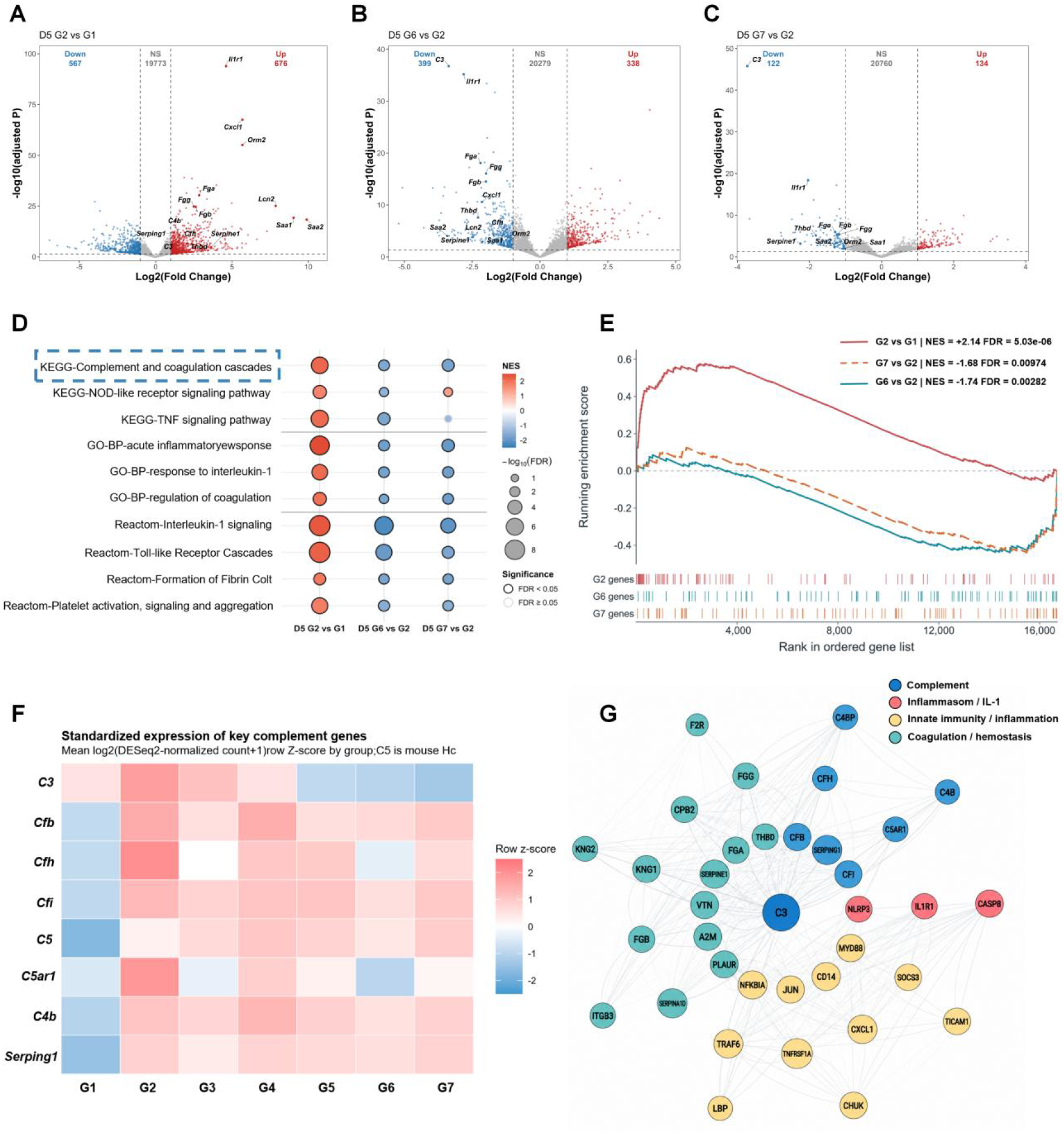
Hepatic transcriptomic analysis reveals alterations in complement-, inflammation-, and coagulation-related pathways following H1N1 infection and combined RG002 and oseltamivir treatment. The experimental groups were defined as follows: G1, healthy control group (Control); G2, H1N1-infected group treated with PBS (PBS); G3, oseltamivir-treated group (OSE); G4, 0.1 mg/kg RG002-treated group; G5, 0.5 mg/kg RG002-treated group; G6, oseltamivir combined with 0.1 mg/kg RG002 treatment group; and G7, oseltamivir combined with 0.5 mg/kg RG002 treatment group. **(A–C)** Volcano plots showing differentially expressed genes in mouse liver tissues on day 5 post-infection. Comparisons include the PBS group versus the Control group **(A)**, the OSE plus 0.1 mg/kg RG002 group versus the PBS group **(B)**, and the OSE plus 0.5 mg/kg RG002 group versus the PBS group **(C)**. Red, blue, and gray dots represent upregulated, downregulated, and non-significantly differentially expressed genes, respectively. DEGs were defined by an adjusted P value < 0.05 and |log_₂_ fold change| ≥ 1. Selected representative genes associated with complement regulation, inflammatory responses, and coagulation-related processes are labeled. **(D)** Bubble plot of gene set enrichment analysis (GSEA) based on the Kyoto Encyclopedia of Genes and Genomes (KEGG), Gene Ontology (GO), and Reactome databases, illustrating the enrichment patterns of complement-, inflammation-, and coagulation-related pathways following H1N1 infection and combination treatment. Bubble colors represent normalized enrichment scores (NES), with red indicating positive enrichment and blue indicating negative enrichment. Bubble sizes correspond to the negative logarithm of the false discovery rate (FDR) [−log_₁₀_(FDR)], while bubble outlines indicate statistical significance (FDR < 0.05). The complement and coagulation cascades pathway is highlighted by a blue dashed box. (E) GSEA enrichment plots of the complement and coagulation cascades pathway, showing pathway enrichment in the PBS group versus the Control group and in the two combination-treatment groups versus the PBS group. (F) Heatmap showing the expression profiles of complement-related genes in liver tissues across the experimental groups. Gene expression values were obtained from DESeq2-normalized counts, log₂-transformed, and subsequently standardized using row-wise Z-scores. Red and blue indicate relatively higher and lower expression levels, respectively. (G) Protein–protein interaction (PPI) network of proteins associated with complement regulation, inflammation, and coagulation. Different colors represent distinct functional modules, including the complement system (blue), inflammasome/IL-1 signaling (pink), innate immunity and inflammatory responses (yellow), and coagulation and hemostasis (teal).

To further characterize the effects of H1N1 infection and therapeutic intervention on hepatic complement-related gene expression, the expression profiles of representative complement-associated genes were examined (Fig. 6F). Compared with the Control group, the PBS group exhibited higher relative expression levels of several complement-related genes, including *C3*, *Cfb*, *Cfh*, *Cfi*, *C5ar1*, *C4b*, and *Serping1*, indicating infection-associated alterations in the transcriptional expression of complement components and regulatory factors. Notably, hepatic *C3* expression was markedly reduced in both the OSE plus 0.1 mg/kg RG002 and OSE plus 0.5 mg/kg RG002 groups compared with the PBS group, consistent with the hepatic C3 mRNA measurements described above. Furthermore, several complement-related genes, including *Cfh*, *Cfi*, and *C5ar1*, exhibited relatively lower expression levels in the OSE plus 0.1 mg/kg RG002 group than in other treatment groups. Collectively, these results indicate that H1N1 infection is associated with alterations in hepatic complement-related gene expression, while combined RG002 and OSE treatment reduces the expression of the RG002 target gene *C3* and is accompanied by selective changes in other complement-related genes, particularly in the lower-dose combination group.

To further explore the potential molecular relationships among complement regulation, inflammatory responses, and coagulation-related processes, a protein–protein interaction (PPI) network was constructed using representative proteins associated with the complement system, inflammasome/IL-1 signaling, innate immunity, and coagulation (Fig. 6G). The resulting network revealed extensive functional associations among these modules. C3 occupied a relatively central position within the network and was closely associated with complement-related proteins, including CFB, CFH, CFI, C4B, and SERPING1. Furthermore, inflammation-related proteins, including IL1R1, NLRP3, CASP8, and MYD88, exhibited functional associations with modules containing coagulation-related proteins, such as FGA, FGB, FGG, and THBD. These findings suggest potential functional interconnections among complement regulation, inflammatory signaling, and coagulation-related biological processes.

## Discussion

In the present study, we investigated the therapeutic potential of RG002, a liver-targeted small interfering RNA (siRNA) targeting complement component C3, in combination with antiviral treatment for alleviating H1N1 influenza virus-associated inflammatory responses and pulmonary tissue injury. Our findings revealed that H1N1 infection was accompanied by increased hepatic C3 mRNA expression, altered serum complement activity, exacerbated inflammatory responses, hematological abnormalities, and pulmonary tissue injury. Further investigations demonstrated that RG002 effectively suppressed hepatic C3 expression in a dose- and time-dependent manner and significantly reduced serum complement-mediated hemolytic activity ex vivo. However, neither RG002 nor oseltamivir (OSE) monotherapy provided limited protection against infection-associated pathological manifestations. In contrast, combined treatment demonstrated greater overall protective effects in alleviating pulmonary inflammation, tissue injury, together with improvements in selected hematological parameters, with OSE plus 0.1 mg/kg RG002 producing the more favorable therapeutic outcomes. These findings suggest that hepatic C3 may represent a potential therapeutic target for simultaneously modulating influenza-associated inflammatory responses, and that combining direct antiviral treatment with appropriately controlled complement modulation may provide more comprehensive protection.

The complement system plays a dual role during influenza virus infection, contributing to antiviral immune defense while potentially mediating inflammation-associated tissue injury (Xiao et al., 2024). During influenza infection, viral components, infection-associated immune complexes, and signals derived from damaged tissues may all contribute to complement activation. Previous studies have demonstrated that complement activation plays an important role in host defense against the 2009 pandemic H1N1 influenza virus, whereas excessive production of complement-derived inflammatory mediators may exacerbate disease progression (Rattan et al., 2017; Wu et al., 2021). Elevated circulating levels of complement activation products, including C3a and C5a, have been reported in patients with H1N1 influenza, suggesting that dysregulated complement activation may contribute to infection-associated inflammatory responses (Ohta et al., 2011). Similar patterns of complement dysregulation have also been observed during severe acute respiratory syndrome coronavirus (SARS-CoV) and SARS-CoV-2 infections. Studies have shown that SARS-CoV infection induces the accumulation of C3 activation products in pulmonary tissues, whereas C3 deficiency attenuates lung injury and inflammatory cell infiltration in infected mice (Gralinski et al., 2018). Furthermore, increased circulating levels of C5a and the terminal complement complex C5b-9 have been observed in some patients with severe COVID-19, accompanied by deposition of complement components, including C4d and C5b-9, within the pulmonary microvasculature. These findings suggest that dysregulated complement activation may contribute to inflammatory responses, vascular endothelial injury, and microvascular pathology associated with severe respiratory viral infections (Java et al., 2020).

Mechanistically, excessive complement activation can promote inflammatory cell recruitment and activation through C3a and C5a, while downstream terminal complement effectors may further amplify inflammatory responses and tissue injury (Detsika et al., 2024). Meanwhile, in certain complement-mediated hematological disorders, deposition of C3 fragments facilitates erythrocyte opsonization and clearance, whereas the membrane attack complex (MAC, C5b-9), can directly mediate erythrocyte membrane damage and lysis (Risitano et al., 2023). These findings suggest that dysregulated complement activation may contribute to inflammatory injury and erythrocyte-associated pathological alterations through distinct downstream effector mechanisms, providing a theoretical rationale for targeting C3, a central component of the complement cascade, to modulate complement-associated inflammatory responses. However, whether these mechanisms contribute to erythrocyte abnormalities during H1N1 infection remains unclear.

Pharmacological intervention studies have provided further experimental evidence supporting the therapeutic potential of complement-targeted strategies for alleviating influenza-associated lung injury. Garcia et al. (2013) demonstrated that inhibition of C5 activation using OmCI significantly reduced neutrophil and macrophage recruitment to the lungs of H1N1-infected mice and attenuated inflammatory responses and pulmonary tissue injury (Garcia et al., 2013). Similarly, Song et al. (2018) reported that blockade of the C5a–C5aR1 signaling pathway alleviated influenza virus-induced acute lung injury and systemic inflammatory responses (Song et al., 2018). Together, these studies provide experimental support for complement-targeted interventions against influenza-associated pulmonary immunopathology. However, these investigations primarily focused on pulmonary inflammation and tissue injury, while the potential of complement-targeted interventions on infection-associated hematological abnormalities remain insufficiently characterized.

Notably, C3 occupies a central position in the complement cascade as the common convergence point of the classical, lectin, and alternative pathways. Its activation contributes not only to the generation of inflammatory effectors, including C3a and C5a, but also under certain pathological conditions to erythrocyte injury through C3b-mediated opsonization and downstream terminal complement activation (Zarantonello et al., 2023). Therefore, compared with therapeutic approaches primarily targeting downstream C5-associated effector mechanisms, targeting C3 may provide broader regulation of the complement cascade at an upstream level, with potential benefits for limiting complement-associated inflammatory responses. Its contribution to H1N1-associated hematological abnormalities, however, requires further investigation.

In the present study, H1N1 infection was accompanied by increased hepatic C3 expression, aggravated pulmonary inflammation, and abnormalities in hematological parameters, suggesting that excessive complement activation may represent a common upstream contributor to these pathological alterations. Given that the liver is the major source of circulating C3, we hypothesized that suppressing hepatic C3 expression to modulate complement function may provide a novel therapeutic strategy for alleviating inflammatory injury associated with H1N1 infection, while hematological alterations were evaluated as additional pathological outcomes.

Our study further validated the C3-targeting activity of RG002 at both the molecular and functional levels. RG002 reduced hepatic C3 expression in a dose- and time-dependent manner, accompanied by decreased serum complement-dependent hemolytic activity, indicating effective target engagement and functional attenuation of circulating complement activity. These findings are consistent with previous studies demonstrating that liver-targeted C3 siRNA can suppress C3 production and ameliorate pathological manifestations in experimental models of complement-associated diseases (Zanchi et al., 2024). Furthermore, we observed that combined RG002 and OSE treatment reduced the levels of IL-1β, IL-6, and IFN-γ in lung tissues during the later stage of H1N1 infection and improved infection-associated alterations in peripheral immune cell composition. Previous studies have demonstrated that C3a can promote NLRP3 inflammasome-associated IL-1β secretion in specific human monocyte models, whereas sublytic MAC formation can also induce NLRP3-dependent IL-1β maturation and release (Asgari et al., 2013; Laudisi et al., 2013). Our hepatic transcriptomic analysis further revealed increased expression of Il1r1 and other inflammation-associated genes following H1N1 infection, accompanied by positive enrichment of IL-1, TNF, and Toll-like receptor-related signaling pathways. In contrast, these pathways generally exhibited negative enrichment following combination treatment. These transcriptomic alterations were broadly consistent with the reduced pulmonary inflammatory cytokine levels, suggesting that modulation of complement-associated inflammatory amplification may contribute to the protective effects of combination therapy. Nevertheless, because the transcriptomic changes were assessed in liver tissue, they cannot directly establish the molecular mechanisms responsible for reduced pulmonary inflammation.

In addition to inflammatory responses, H1N1 infection was associated with hematological abnormalities, including decreased red blood cell counts and hemoglobin levels, accompanied by elevated lactate dehydrogenase (LDH) and altered bilirubin levels during the early stage of infection. Under certain pathological conditions, excessive complement activation can induce erythrocyte membrane damage through the formation of the terminal complement complex, MAC (Risitano et al., 2023). Clinical evidence further supports the therapeutic value of upstream C3 inhibition, as pegcetacoplan has been shown to improve hemoglobin levels and hemolysis-related outcomes in patients with paroxysmal nocturnal hemoglobinuria (Hillmen et al., 2021). In the present study, RG002 reduced serum complement-mediated hemolytic activity ex vivo using serum from uninfected treated mice, while combination treatment improved selected hematological parameters and biochemical abnormalities, including LDH and bilirubin levels, suggesting that complement modulation may contribute to the alleviation of infection-associated erythrocyte abnormalities. Beyond erythrocyte injury, increasing evidence indicates extensive bidirectional crosstalk between the complement and coagulation systems. In an experimental model, evidence demonstrated that inhibition of C3, C5, or C5aR1 attenuated monocyte tissue factor expression and coagulation activation (Gravastrand et al., 2019), supporting a mechanistic link between complement activation and procoagulant responses. Clinical evidence further supports this relationship, as treatment with the C5 inhibitor eculizumab has been associated with a marked reduction in thromboembolic events in patients with paroxysmal nocturnal hemoglobinuria (Hillmen et al., 2007). Consistent with this concept, our hepatic transcriptomic analysis revealed positive enrichment of the complement and coagulation cascades following infection, accompanied by increased expression of fibrinogen-related genes, including *Fga, Fgb,* and *Fgg*. In contrast, these pathways exhibited negative enrichment following combination treatment. These findings suggest that complement-directed immunomodulation, together with antiviral therapy, may influence the hepatic inflammatory and coagulation-associated responses to H1N1 infection. However, LDH and bilirubin are not specific markers of hemolysis, and changes in fibrinogen-related gene expression do not directly establish coagulation dysfunction. Therefore, further studies are required to determine whether hepatic C3 suppression directly modulates complement-associated erythrocyte injury and coagulation abnormalities during H1N1 infection.

Collectively, these findings suggest that modulating systemic complement function through hepatic C3 targeting may alleviate infection-associated inflammatory injury, accompanied by improvements in selected hematological abnormalities and biochemical parameters. These observations further support C3 as a potential therapeutic target for simultaneously addressing these pathological manifestations of H1N1 infection.

Despite effectively modulating complement activity, RG002 monotherapy provided limited protection, which may be related to the dual role of complement in influenza virus infection. Kopf et al. (2002) demonstrated that C3 deficiency impaired virus-specific T-cell priming and recruitment to the lungs, resulting in delayed viral clearance (Kopf et al., 2002). Similarly, O’Brien et al. (2011) confirmed the protective role of C3 in H1N1 and H5N1 influenza infection models (O’Brien et al., 2011). Notably, in our study, RG002 monotherapy at 0.5 mg/kg did not confer substantial overall protection and was associated with relatively higher pulmonary viral RNA level at certain time points, suggesting that stronger C3 suppression may adversely affect viral control. In addition, although OSE limits viral spread by inhibiting influenza virus neuraminidase (Zhang et al., 2025), antiviral monotherapy may be insufficient to fully control host immunopathological responses already initiated during infection. Given their distinct mechanisms of action, combining these two treatments may provide complementary benefits by reducing viral stimulation and modulating complement-associated inflammatory responses (Pandey et al., 2023). In the present study, OSE combined with 0.1 mg/kg RG002 demonstrated more favorable overall therapeutic outcomes than the higher-dose combination. Hematoxylin and eosin (H&E) and periodic acid–Schiff (PAS) staining further revealed more pronounced improvements in pulmonary inflammation and histopathological alterations. Although pulmonary viral RNA levels tended to be lower in the combination treatment group than in the OSE monotherapy group, the difference did not reach statistical significance. These observations suggests that the additional protective effects of combination therapy may be associated more closely with the alleviation of host pathological responses than with enhanced suppression of viral replication alone. Taken together, these findings indicate that the relationship between the extent of C3 suppression and therapeutic efficacy may not be linear, and that appropriately controlled complement modulation, rather than maximal C3 suppression, may be more beneficial for balancing antiviral immunity and tissue protection.

This study provides experimental evidence supporting the potential of liver-targeted C3 silencing combined with antiviral treatment against H1N1 infection. Nevertheless, several limitations should be acknowledged. First, although changes in C3 expression and serum complement activity were detected, complement activation products, including C3a, C5a, and C5b-9, as well as local complement deposition in lung tissues, were not systematically evaluated. Therefore, the specific downstream mechanisms by which RG002 modulates complement-associated inflammatory injury remain unclear. The absence of a non-targeting siRNA control (siNC) also limits our ability to attribute the observed effects specifically to C3 knockdown. Second, although improvements in erythrocyte-associated hematological parameters and biochemical abnormalities were observed, direct evidence, including erythrocyte surface complement deposition, plasma-free hemoglobin, and haptoglobin measurements, is still lacking. Consequently, the mechanisms underlying infection-associated hematological abnormalities remain to be established. Third, hepatic transcriptomic data and some pulmonary and hematological measurements were obtained from different animals or experimental batches, preventing the establishment of individual-level causal relationships between molecular alterations and phenotypic outcomes. Furthermore, hepatic transcriptional changes may not fully reflect local pulmonary inflammatory signaling, and the principal transcriptomic comparisons did not directly establish the additional molecular effects of RG002 beyond OSE monotherapy. In addition, sample loss due to animal mortality during the later stages of infection may introduce survivor bias, while the limited number of RG002 dose levels precludes identification of an optimal therapeutic range. Future studies should further investigate the extent of complement suppression, viral clearance kinetics, and immunological safety across different RG002 doses and administration schedules, as well as the independent therapeutic contribution of RG002 when added to OSE treatment.

In conclusion, our findings suggest that appropriately controlled complement modulation through liver-targeted C3 silencing, combined with direct antiviral treatment, may provide a dual therapeutic strategy for alleviating inflammatory responses and pulmonary tissue injury. This approach offers a potential new direction for host-targeted adjunctive therapy against influenza and other severe respiratory viral infections.

## Materials and Methods

### Animals and Ethical Statement

All mice used in this study were female BALB/c mice (6–8 weeks old), purchased from GemPharmatech Co.,Ltd. and SPF (Beijing) Biotechnology Co., Ltd. All mouse experiments were conducted in accordance with the Guidelines for the Humane Care and Use of Laboratory Animals and were approved by the Institutional Animal Care and Use Committee (IACUC) of Chinese Center for Disease Control and Prevention (Approval No.: bdbs20250422054). All protocols strictly followed relevant guidelines and policies.

Mice were housed in a specific pathogen-free (SPF) facility under controlled environment with a constant temperature of 25°C, relative humidity of 50%, and a 12-hour light/dark cycle. Sterile water and food were provided ad libitum throughout the study. A humane endpoint was defined as a loss of 25% or more of baseline body weight.. Mice reaching this endpoint were immediately and humanely euthanized in strict accordance with institutional protocols.

### Experimental Materials

Influenza A virus (H1N1, PR8 strain) was obtained from the Institute for Viral Disease Control and Prevention, Chinese Center for Disease Control and Prevention. The virus was amplified in 9-11-day-old specific-pathogen-free (SPF) chicken embryos, aliquoted and stored at −80 °C for subsequent use. Viral titers were determined by plaque assay using MDCK cells. The 50% lethal dose (LD₅₀) of this virus in mice was pre-determined to be 1.5 × 10⁸ PFU/mouse.

Candidate drug RG002 was synthesized and provided by Suzhou Xuanjing Biotechnology Co., Ltd. Before use, 20 mg of RG002 powder was dissolved in 1 mL of 0.9% sodium chloride solution to prepare a 20 mg/mL stock solution, which was aliquoted and stored at −20 °C. Oseltamivir phosphate (OSE, Cat. No. HY-17016) was purchased from MedChemExpress (MCE, Shanghai, China). Briefly, 1 g of oseltamivir phosphate was dissolved in 25 mL of 0.9% sodium chloride solution, with sonication to aid dissolution, yielding a 40 mg/mL stock solution. Assuming an average mouse body weight of 20 g, the stock solution was further diluted to a 5 mg/mL working solution, and 200 μL of the working solution was administered to each mouse via intragastric gavage.

Isoflurane (Cat. No. R510-220) was obtained from RWD Life Science Co., Ltd.(Shenzhen, China). Protease inhibitor (Cat. No. A8260) was purchased from Solarbio (Beijing, China). The nucleic acid extraction and purification kit for viral load detection (Cat. No. T332) was acquired from Tianlong Science and Technology Co., Ltd.(Xi’an, China). The One Step PrimeScript RT-PCR Kit (Cat. No. RR064A) was purchased from Takara Biomedical Technology Co., Ltd.,(Beijing, China).

RNA isolater Total RNA Extraction Reagent (Cat. No. R401-01), used for detection of complement C3 mRNA expression and hepatic transcriptome sequencing, was obtained from Vazyme Biotech Co., Ltd. (Nanjing, China). The 5× Hifair One Step RT SuperMix (Cat. No. 11142ES) and Hieff qPCR SYBR Green Master Mix (Cat. No. 11201ES) were both purchased from Yeasen Biotechnology Co., Ltd.(Shanghai, China).

Mouse IL-6 and IFN-γ ELISA kits (Cat. Nos. ED-20188, ED-20140) were supplied by LCS Biotechnology Co., Ltd.(Xiamen, China). Mouse IL-1 β ELISA kits (Cat No. BY-EM220174) was obtained from BYabscience technology Co., Ltd. (Nanjing, China).

Mouse complement C3 protein ELISA kit (Cat. No. ab157711) was obtained from Abcam, UK. Mouse complement alternative pathway ELISA kit (Cat. No. HIT422) was purchased from HycultBiotech, the Netherlands. Sheep red blood cells (SRBC, Cat. No. HQ80073-002) and rabbit anti-sheep red blood cell antiserum (hemolysin, Cat. No. HQ40011) were purchased from Hongquan Biotechnology Co., Ltd.,(Guangzhou, China).

### H1N1 Infection Mouse Model

To characterize inflammatory responses and complement-pathway activation following H1N1 infection, BALB/c mice (6-8 weeks old) were randomly assigned to different groups. Mice in the infection group were anesthetized, then intranasally inoculated with H1N1 influenza virus at a dose of 3 LD₅₀ (50 μL per mouse). The control group received an equal volume of PBS via intranasal instillation. Lung tissues were harvested on days 1, 3, 5, and 7 post-infection for viral load measurement, cytokine quantification and histopathological evaluation. Liver tissues were collected for detection of complement C3 mRNA expression and transcriptome sequencing. Peripheral blood was obtained to analyze complement C3 concentration, complement alternative-pathway activity, and complete blood count (CBC) and serum biochemical analysis.

### Pharmacokinetics and in-vitro Hemolysis Assay of RG002

To evaluate the pharmacokinetic profiles of RG002, naïve mice were randomly divided into a blank control group and four treatment groups, which received subcutaneous injection of RG002 at doses of 0.1 mg/kg, 0.5 mg/kg, 1 mg/kg and 3 mg/kg, respectively. Liver tissues were collected at five predefined sampling time-points within 20 days post-administration (Day 1, 2, 6, 13, and 20). Real-time quantitative PCR (RT-qPCR) was used to detect changes in hepatic complement C3 mRNA levels. Meanwhile, serum samples from the 0.1 mg/kg, 0.5 mg/kg and 1 mg/kg dosage groups were harvested for the in-vitro hemolysis assay. A separate survival subgroup was established for continuous recording of body weight changes.

### In-vivo Therapeutic-Efficacy Evaluation

To assess the in-vivo therapeutic effects of RG002 monotherapy at different doses, as well as combination therapy of RG002 and oseltamivir phosphate (OSE), mice were randomly divided into seven groups (n = 6 per group):

1. Healthy control group;
2. Infected PBS vehicle group;
3. Positive-control oseltamivir group;
4. 0.1 mg/kg RG002 monotherapy group;
5. 0.5 mg/kg RG002 monotherapy group;
6. OSE + 0.1 mg/kg RG002 combination-therapy group;
7. OSE + 0.5 mg/kg RG002 combination-therapy group.

Except for the healthy control group, mice were intranasally challenged with H1N1 virus at 3 LD₅₀ on day 0. Therapeutic interventions were initiated immediately after infection. Mice in the PBS vehicle group, RG002 monotherapy groups and RG002-OSE combination groups received a single subcutaneous injection of PBS or RG002 on the day of infection. Mice in the OSE monotherapy group and OSE-RG002 combination groups were administered OSE by intragastric gavage once daily starting on the day of infection until the experimental endpoint or human endpoint. RG002 was administered as a single subcutaneous bolus, whereas oseltamivir was given by daily gavage for sustained therapeutic treatment. Mouse survival rates and body-weight changes were recorded daily.

An independent parallel cohort of mice was set up for scheduled sample collection. Blood, liver, and lung tissues were collected on days 1, 3, 5 and 7 post-infection for subsequent measurements of viral load, inflammatory cytokines, complement C3 levels, serum biochemistry and histopathological analysis.

### Viral Load Detection

Mouse lung tissues were weighed and thoroughly homogenized in pre-chilled PBS at a ratio of 4 volumes of PBS per gram tissue (1 g tissue : 4 mL PBS) using a chilled tissue homogenizer. After centrifugation, supernatants were harvested. Total RNA was extracted with the Tianlong T332 nucleic acid extraction kit on an automated nucleic acid extractor (Tianlong). Viral nucleic acids were subjected to reverse-transcription quantitative PCR (RT-qPCR) amplification using the One Step PrimeScript RT-PCR Kit (TaKaRa), and signals were detected on a LightCycler 96 real-time PCR system (Roche).

The PCR thermal profile was set as follows: 37 °C for 120 s, initial denaturation at 95 °C for 30 s, followed by 45 cycles of 95 °C for 10 s and 60 °C for 30 s. Primers were designed targeting the relatively conserved region of the matrix protein (M) gene of influenza A virus. The primer sequences are shown below:

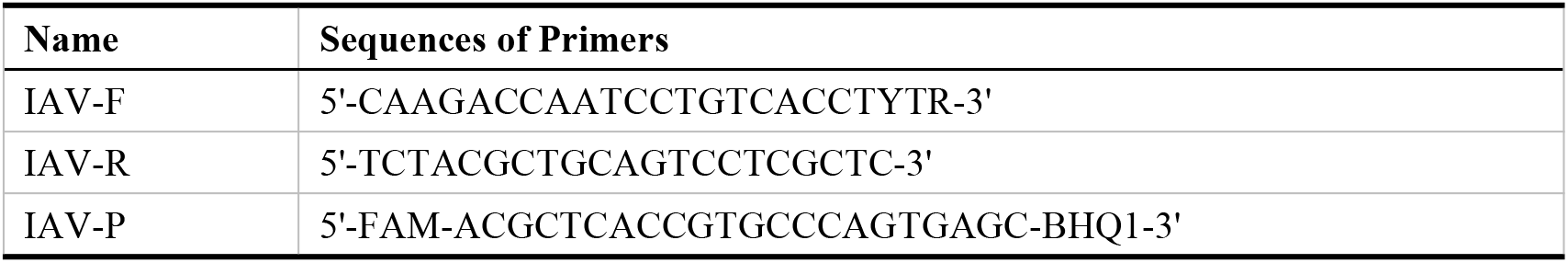

### Analysis of Complement C3 mRNA Expression

Mouse liver tissues were collected and thoroughly homogenized with 500 μL TRIzol reagent using a chilled tissue homogenizer. Total RNA was extracted via the classic phenol-chloroform method. Briefly, chloroform was added to the homogenate, followed by vigorous vortexing for phase separation. After centrifugation, the aqueous phase was transferred to a new tube. RNA was precipitated with isopropanol. The RNA pellet was washed with 75% ethanol, air-dried at room temperature, and resuspended in RNase-free ultrapure water. RNA concentration and purity were measured using a NanoDrop spectrophotometer.

Purified total RNA was reverse-transcribed into cDNA using reverse-transcription Kit (Yeasen Biotechnology). Quantitative real-time PCR (qPCR) was subsequently performed with qPCR master mix (Yeasen Biotechnology) on a QuantStudio 3 Real-Time PCR System (Thermo Fisher Scientific, USA). GAPDH was applied as the reference gene. Relative mRNA expression of the hepatic C3 gene was calculated using the 2−ΔΔCt method after normalization against the reference gene. The primer sequences are listed below:

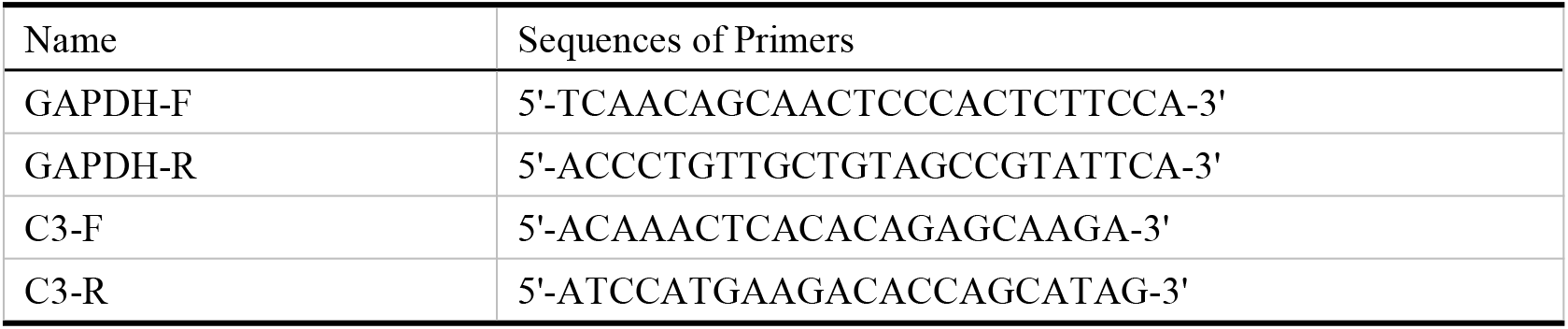

### Analysis of Complement C3 Protein and Complement Alternative-Pathway Activity

Collected mouse blood was incubated at 4 °C for more than 30 min, followed by centrifugation at 2500 rpm for 15 min to separate serum from blood cells. The supernatant was harvested and subjected to a second centrifugation at 2500 rpm for 10 min to completely remove residual blood cells. Serum complement C3 protein concentrations were detected using an ELISA kit (Abcam), and serum complement alternative-pathway activity was measured with an ELISA kit (Hycult Biotech). Absorbance was read at 450 nm on a multimode microplate reader (Cytation 5, BioTek, USA), and the complement C3 actual concentration and alternative pathy activity of each analyte were calculated accordingly.

### Cytokine Analysis

Mouse lung tissues were weighed and mixed with PBS supplemented with protease inhibitors at a 1:9 weight-to-volume ratio. Tissues were fully homogenized using a pre-chilled tissue homogenizer. The homogenate was centrifuged at 5000 rpm for 10 min, and the supernatant was collected. The supernatant was diluted 15-fold with 1×PBS (supernatant:PBS = 1:14). Secretion levels of IL-6, IFN-γ, and IL-1β were quantified using corresponding ELISA kits. Absorbance values were acquired at 450 nm with a multimode microplate reader (Cytation 5, BioTek, USA). Actual analyte concentrations were calculated based on the standard curve generated from kit reference standards.

### In-vitro Serum Hemolysis Assay

The hemolytic activity of serum from RG002-treated mice was evaluated using a modified complement-dependent hemolysis assay adapted from the classical CH50 protocol, in which hemolysis percentage rather than CH50 titer was determined. First, hemolysin was diluted 1:100 with 1×PBS; briefly, 20 μL hemolysin was mixed with 1980 μL PBS to obtain a total volume of 2 mL for sensitizing sheep red blood cells (SRBC). An equal volume of 2% SRBC suspension was mixed with the diluted hemolysin (2 mL 2% SRBC plus 2 mL diluted hemolysin, total volume = 4 mL). The mixture was incubated for 30 min at 37 °C in an Eppendorf thermal block protected from light to yield sensitized SRBC suspension with a final cell concentration of 1%.

Aliquots (90μL each) of serum from RG002-treated mice (0.1 mg/kg, 0.5 mg/kg, 1.0 mg/kg groups) and naïve mouse serum were mixed with sensitized SRBC suspension (30μL each) and co-incubated at 37 °C in a thermal block for 30 min. Samples were centrifuged at 2000 rpm for 10 min. The supernatants were transferred to a 96-well plate (100μL per well), and absorbance was measured at 542 nm using a microplate reader.

Four reference hemolysis control tubes were prepared in parallel: 100% hemolysis control, 50% hemolysis control, 25% hemolysis control, and no-hemolysis control. The 100% hemolysis control supernatant was obtained by mixing 2 mL of 2% SRBC suspension with 14 mL double-distilled water (ddH₂O), followed by centrifugation. The 50% hemolysis control was prepared by mixing the 100% hemolysis supernatant with ddH₂O at a 1:1 volume ratio. The 25% hemolysis control was generated by mixing the 50% hemolysis supernatant with ddH₂O at a 1:1 volume ratio. The no-hemolysis control was prepared by mixing 1 mL SRBC suspension with 4 mL PBS. All experimental, 100% hemolysis control, and o-hemolysis control reactions were adjusted to maintain a final SRBC concentration of 0.25%.

The hemolysis rate (%) for each treatment group was calculated using the following formula:

Hemolysis rate (%)= (ODexperimental-ODbackground)/(OD100% hemolysis-ODbackground) ×100% where ODexperimental is the absorbance of the sample group, ODbackground is the absorbance of the none-hemolysis control, and OD100% hemolysis is the absorbance of the complete-hemolysis control.

### Complete Blood Count and Serum Biochemical Assays

At the experimental endpoint of the pharmacodynamic study, blood samples were collected from mice in each group. Aliquots of whole blood and isolated serum were submitted to Wuhan Seville Biotechnology Co., Ltd. (ServiceBio) for detection. Hematological parameters were measured using a Mindray veterinary automated hematology analyzer (BC-2800vet). Concentrations of total bilirubin (TBIL), direct bilirubin (DBIL) were determined by the vanadate oxidation method, and indirect bilirubin (IBIL) as the difference between TBIL and DBIL. Serum lactate dehydrogenase (LDH) activity was determined using the lactate substrate method.

### Histopathological Examination

Mouse lung tissues were harvested and fixed in 4% paraformaldehyde for several hours, followed by gradient ethanol dehydration and paraffin embedding. Tissues were cut into serial sections. After deparaffinization and rehydration, sections were subjected to hematoxylin-eosin (H&E) staining and periodic-acid-Schiff (PAS) staining, respectively. Pathological alterations were observed and images were captured under a light microscope. H&E-stained sections were used to evaluate the extent of inflammatory cell infiltration and structural damage in lung tissue. PAS staining was applied to assess airway mucus secretion and airway epithelial injury.

Inflammatory cell infiltration was quantitatively assessed using H&E-stained lung tissue sections. Five randomly selected, non-overlapping fields of the lung parenchyma were imaged for each mouse using the same magnification and identical image acquisition settings. Image analysis was performed using ImageJ software. Areas containing densely clustered, intensely hematoxylin-stained nuclei were identified by color thresholding and defined as inflammatory cell-rich regions. The same threshold parameters were applied to all images within the same experiment. The areas of the inflammatory cell-rich regions and the total evaluable lung tissue area were measured in each field. The percentage of inflammatory infiltration area was calculated as follows:

Inflammatory infiltration area (%) = inflammatory cell-rich area /total evaluable lung tissue area × 100%.

PAS-stained lung sections were further used to quantify airway epithelial injury. Five non-overlapping fields containing clearly identifiable airway structures were selected from each mouse for evaluation. Airway epithelial injury was defined as disruption or detachment of the epithelial lining, marked morphological abnormalities or disorganization of epithelial cells, or focal exposure of the basement membrane. The lengths of the injured airway basement membrane and the total airway basement membrane perimeter were measured using ImageJ software. The proportion of airway epithelial injury was calculated as follows:

Airway epithelial injury (%) = length of injured airway basement membrane/total airway basement membrane perimeter × 100%.

The mean value obtained from the five fields was calculated for each mouse and used as one biological replicate in subsequent statistical analyses.

### Hepatic Transcriptome Sequencing and Differential Gene Expression Analysis

Total RNA was extracted from mouse liver tissue homogenates as described above. Following RNA quality assessment, library construction and high-throughput transcriptome sequencing were performed by Shenzhen BGI Technology Co.,Ltd. (Shenzhen, China).Eukaryotic mRNA libraries were sequenced on the DNBSEQ platform using a paired-end 150-bp (PE150) strategy. Raw sequencing reads were subjected to quality filtering by the sequencing service provider, and the resulting clean reads were provided for subsequent bioinformatic analyses. Clean reads were aligned to the mouse reference genome (GRCm39), followed by gene expression quantification, normalization, and differential expression analysis between experimental groups. Differentially expressed genes (DEGs) were identified based on adjusted P values and fold changes, and gene expression differences between groups were visualized using volcano plots.

To further characterize the biological processes associated with H1N1 infection and therapeutic intervention, gene set enrichment analysis (GSEA) was performed using clusterProfiler (v4.18.4) with the fgsea backend (v1.36.2). All testable and successfully mapped genes were ranked according to the Wald statistics obtained from DESeq2. Gene sets from Gene Ontology Biological Process (GO-BP), Kyoto Encyclopedia of Genes and Genomes (KEGG), and Reactome were used for functional enrichment analysis. GO-BP annotations were obtained from org.Mm.eg.db (v3.22.0), KEGG gene sets from the online mouse KEGG database, and Reactome gene sets from MSigDB 2026.1. Reactome gene sets were mapped to mouse orthologs using msigdbr (v26.1.1). Gene set sizes were restricted to 10–500 genes for GO-BP and KEGG analyses and 5–1,000 genes for Reactome analysis. Statistical significance was defined as an adjusted P value or q value < 0.05, as specified in the corresponding figure legends.

For heatmap visualization, normalized expression values of representative complement-related genes were transformed using log₂(DESeq2-normalized counts + 1). Gene-wise Z-scores were calculated across all samples, and group-level mean Z-scores were used to visualize and compare gene expression patterns among experimental groups.

Additionally, representative proteins associated with complement activation, inflammation, and coagulation were selected to construct a protein–protein interaction (PPI) network. These proteins were further classified into functional modules according to their biological roles to explore potential functional associations among these processes.

### Statistical Analysis

Statistical analyses of non-transcriptomic outcomes were performed using GraphPad Prism (version 9.5.0). Data are presented as mean ± SD. A two-tailed unpaired Student’s t-test was used for comparisons between two groups. One-way analysis of variance (one-way ANOVA) followed by Tukey’s multiple-comparisons test was adopted for comparisons among multiple groups at a single time point. For continuous variables covering multiple time points and different treatment groups, two-way analysis of variance (two-way ANOVA) coupled with Sidak’s multiple-comparisons test was applied for multiple-comparison analysis. Mouse survival curves were plotted by Kaplan–Meier survival analysis, and intergroup differences in survival were assessed by the Log-rank (Mantel-Cox) test. A Pvalue < 0.05 was considered statistically significant

## Compliance and ethics

All animal experimental procedures were approved by the Institutional Animal Care and Use Committee (IACUC) of China CDC (Approval No. bdb620250422054) and conducted in accordance with the institutional guidelines for the care and use of laboratory animals.

## Author Contributions

Gege Shi and Zixin Tu contributed equally to this work. Gege Shi and Zixin Tu performed the experiments, analyzed and interpreted the data, and drafted the manuscript. Yu Li, Yu Ou, Yizheng Ao, Haoyuan Feng, and Xiaofeng Han contributed to experimental investigation and data collection. Yawei Wang, Shangqing Gao, Qi Ma, Xinyu Gao, and Zimu Li contributed to experimental procedures, data acquisition, and analysis. Haitao Li and Minghui Yang provided technical support and contributed to data interpretation. Xing-Jie Liang and Jinchao Zhang provided resources and critical scientific input. Baoying Huang, Yuanyu Huang, and Yuhua Weng contributed to conceptualization, supervision, and project administration. Baoying Huang, Yuanyu Huang, and Yuhua Weng critically revised the manuscript. All authors read and approved the final manuscript.

## Data availability

The original contributions presented in this study are included in this article. Further inquiries can be directed to the corresponding authors.

