## Supplementary Figures for "C3-targeting siRNA Combined with Oseltamivir Alleviates Inflammation and Lung Injury in H1N1-Infected Mice"

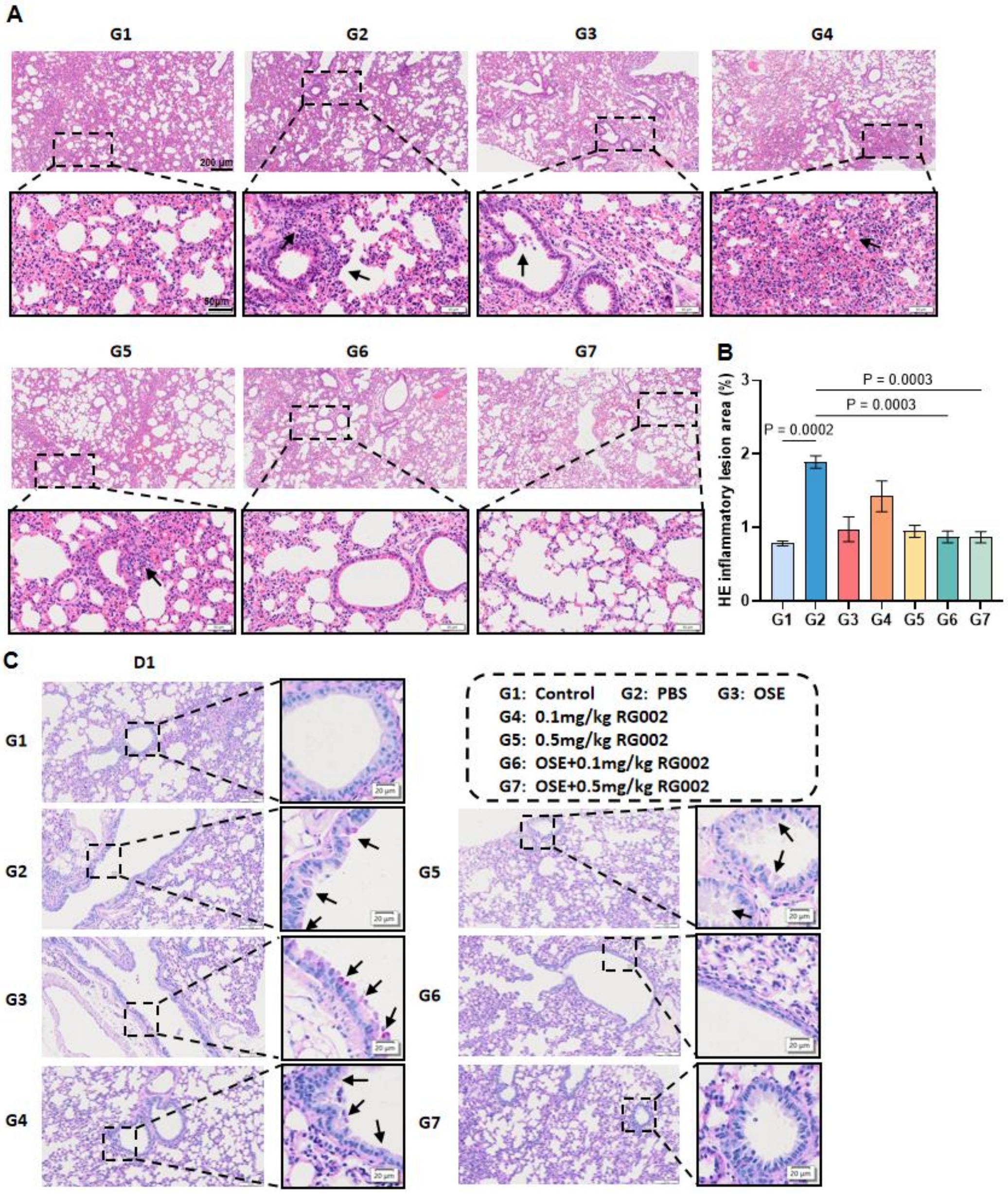


**Supplementary Figure 1. RG002 combined with OSE alleviates pulmonary inflammation and airway epithelial injury in H1N1-infected mice on Day 1. (A)** Representative H&E-stained lung sections on Day 1 after infection, showing inflammatory-cell infiltration and structural alterations in the lung tissues of the different treatment groups. Black arrows indicate areas of inflammatory-cell accumulation or tissue injury. Scale bars, 200 μm in the main images and 50 μm in the enlarged images. **(B)** Quantitative analysis of inflammatory lesion areas in H&E-stained lung sections on Day 1. **(C)** Representative PAS-stained lung sections on Day 1 showing airway epithelial integrity, epithelial cell detachment, and PAS-positive material deposition in the different treatment groups. Black arrows indicate PAS-positive material and detached airway epithelial cells. Scale bars, 100 μm in the main images and 20 μm in the enlarged images. Data are presented as mean ± SD， n = 3 mice per group. Statistical comparisons were performed using ordinary one-way ANOVA.
